# Zebrafish spinal cord repair is accompanied by transient tissue stiffening

**DOI:** 10.1101/666032

**Authors:** Stephanie Möllmert, Maria A. Kharlamova, Tobias Hoche, Anna V. Taubenberger, Shada Abuhattum, Veronika Kuscha, Thomas Kurth, Michael Brand, Jochen Guck

## Abstract

Severe injury to the mammalian spinal cord results in permanent loss of function due to the formation of a glial-fibrotic scar. Both the chemical composition and the mechanical properties of the scar tissue have been implicated to inhibit neuronal regrowth and functional recovery. By contrast, adult zebrafish are able to repair spinal cord tissue and restore motor function after complete spinal cord transection owing to a complex cellular response that includes neurogenesis and axon regrowth. The mechanical mechanisms contributing to successful spinal cord repair in adult zebrafish are, however, currently unknown. Here, we employ AFM-enabled nano-indentation to determine the spatial distributions of apparent elastic moduli of living spinal cord tissue sections obtained from uninjured zebrafish and at distinct time points after complete spinal cord transection. In uninjured specimens, spinal gray matter regions were stiffer than white matter regions. During regeneration after transection, the spinal cord tissues displayed a significant increase of the respective apparent elastic moduli that transiently obliterated the mechanical difference between the two types of matter, before returning to baseline values after completion of repair. Tissue stiffness correlated variably with cell number density, oligodendrocyte interconnectivity, axonal orientation, and vascularization. The presented work constitutes the first quantitative mapping of the spatio-temporal changes of spinal cord tissue stiffness in regenerating adult zebrafish and provides the tissue mechanical basis for future studies into the role of mechanosensing in spinal cord repair.

## Introduction

The spinal cord contains neurons and glia that act collectively to transmit sensory information from the peripheral nervous system to the brain, and motor commands from the brain to the periphery of the body evoking both voluntary and involuntary movements. After traumatic spinal cord injury in mammals, this information exchange is irreversibly impaired due to the immediate disruption of axonal projections, neuronal cell death and the eventual formation of a glial-fibrotic scar [1–4]. The scar tissue not only inhibits axonal regrowth across the lesion site due to its biochemical composition [4], but has also been proposed to act as a mechanical impediment [2]. Analogous to SCI in mammalian systems, traumatic spinal cord injury in zebrafish entails the immediate loss of function caudal to the lesion site [5, 6]. However, the absence of voluntary body movements caudal to the injury level is, in contrast to mammalian paralysis, not permanent [5, 6]. Zebrafish respond to spinal cord injury by a complex cellular response including proliferation [7–9], migration [10], differentiation [8, 9, 11] and morphological changes [8, 9]. New motor neurons originating from proliferating radial glial cells mature and eventually form synaptic connections [8]. Severed axons that descend from the brainstem regrow, traverse the injury site and innervate the caudal spinal cord [7, 12]. Fibroblast-like cells accumulate in the injury site, secrete collagen XII and thereby contribute to an ECM that is growth promoting for axons [13]. These processes restore the spinal cord tissue and facilitate functional recovery in adult zebrafish within 6-8 weeks post-injury [8].

Morphological changes, proliferation, migration and differentiation also constitute responses that mechanosensitive neurons and glia exhibit when exposed to distinct mechanical environments [14–17]. *In vitro* studies of neural cells reported an increased branching of neurons on compliant, but directed axonal growth on stiff substrates [14, 18]. Astrocytes and microglia display morphological characteristics of an activated phenotype and upregulate inflammatory genes and proteins when exposed to a mechanical stimulus that deviates from their physiological mechanical environment both *in vitro* and *in vivo* [17]. Oligodendrocyte precursor cells increase their expression of myelin basic protein and display an elaborated myelin membrane on stiffer substrates as compared to more compliant substrates indicating a preferred mechanical environment for differentiation [15].

*In vivo*, this mechanical environment is formed by the surrounding nervous tissue whose mechanical properties are determined by factors such as the combined material properties of neighboring cells, cell density, myelin content, collagen content, extra cellular matrix composition and cell interconnectivity [19, 20]. As these may change during development or after pathological events, concomitant changes of mechanical tissue properties and their direct involvement in a wide range of CNS conditions and diseases becomes apparent [21–24]. Axonal growth during optic tract development in *Xenopus laevis*, for instance, is guided by temporally changing stiffness gradients in adjacent brain tissue [18]. Acute demyelination in mouse models of multiple sclerosis is accompanied by an increase of stiffness in affected brain regions that were hypothesized to present a mechanically subideal environment to support potentially remyelinating oligodendrocytes [25].

The aforementioned examples of neural mechanosensitivity and stiffness changes of nervous tissues during developmental and pathological events suggest an intricate interplay between neural cell types and the mechanical properties of the nervous tissues in which they reside. Neurogenesis, axonogenesis and the resultant functional recovery after spinal cord injury in adult zebrafish might be likewise accompanied by, or even causally linked to the mechanical changes of the spinal parenchyma. Here, we have characterized the mechanical properties of the adult zebrafish spinal cord. AFM-enabled indentation measurements of acutely prepared living spinal cord slices revealed that spinal cord tissues display a homeostatic mechanical phenotype in which gray matter is stiffer than white matter along the anterior-posterior axis. The re-establishment of tissue homeostasis after complete spinal cord transection was accompanied by transient tissue stiffening. Hence, the cell types and processes associated with functional recovery in adult zebrafish are exposed to spatio-temporally changing mechanical signals provided by the surrounding spinal parenchyma in adult zebrafish and could thus be influenced by these changing mechanical cues. Interestingly, the successful regrowth of axons across the lesion site was associated with an *increase* in the stiffness of the tissue, quite opposite of what might have been expected if a stiff glial scar constituted a mechanical barrier. This finding forms a solid, quantitative foundation for further studies into the causal relationship between mechanosensing and functional repair.

## Methods

All animal experiments were conducted according to the guidelines of the German Animal Welfare Act and under the supervision of the Regierungspräsidium Dresden (DD24.1-5131/339/5 and D24-5131/338/52).

### Zebrafish lines

All zebrafish were kept and bred under standard conditions as described in [26]. The transgenic line Tg(mbp:GFP) was established and provided by the laboratories of Cheol-Hee Kim, Chungnam National University, South Korea, and Hae-Chul Park, Korea University Ansan Hospital, South Korea [27]. The transgenic line Tg(alpha1-tubulin:mls-dsRed) was established in the laboratory of Carla Koehler, UCLA, USA and provided by Christopher Antos, CRTD, Germany. All experiments were carried out with Tg(mbp:GFP, alpha1-tubulin:mls-dsRed) fish and wild type fish (*wik*). All experiments comprise male and female fish.

### Spinal cord dissection for indentation measurements

All zebrafish were sacrificed by immersion in ethyl 3-aminobenzoate methanesulfonate (MS-222, 0.1% in PBS, Sigma-Aldrich, A5040) until five minutes after the respiratory movement of the opercula stopped. This was followed by subsequent immersion in ice-cold water as recommended in [28]. Sacrificed zebrafish were pinned to a silicone-covered petri dish and placed under a stereo microscope. First, dorsal scales were scrapped off. A scalpel was then used to transversely incise the muscle tissue near the brain stem and moved caudally with a sawing motion. To expose the vertebral column, forceps with different tip dimensions were used to remove remaining muscle tissue and carefully break away extending spinal processes. Once the spinal cord was fully exposed, thin forceps were used to gently detach the meninges. It has proven beneficial to keep the pia mater intact as its removal can introduce structural damage to the spinal cord and impede accurate vibratome sectioning. The desired length of tissue was separated from the remaining spinal cord by two incisions and levered out of the vertebral column using the tips of closed forceps. To wash away remaining blood and adipose cells, the severed spinal cord piece was placed in cold artificial cerebrospinal fluid (aCSF) that contained (in mM) 134 NaCl, 2.9 KCl, 1.2 MgCl_2_, 2.1 CaCl_2_, 10 HEPES buffer, and 10 glucose, adjusted to pH 7.8 with NaOH [29]. At last, laterally extending nerve fibers were abscised along the dissected spinal cord tissue as they can lead to the tissue being pulled out of the agarose embedding during vibratome sectioning. To indicate the tissue’s directionality after dissection, various strategies were employed. For instance, the posterior part of the medulla oblongata, whose diameter is distinctly greater than that of the spinal cord, was exposed and excised along with the spinal cord tissue. Melanophores that cover the spinal cord’s surface were left untouched and used for orientation. The caudally decreasing diameter of the spinal cord served as an additional indicator of directionality, but this evaluation required careful visual inspection and has proven suitable only for the dissection of great lengths of spinal cord tissue or entire spinal cords.

### Tissue embedding and vibratome sectioning for indentation measurements

Dissected spinal cord tissue was immersed in liquid low-gelling-point agarose (2.5% in aCSF, cooled to 31°C, Sigma-Aldrich, A0701). The tissue was then centered and straightened with insect pins. Upon solidification, a piece of agarose gel containing the spinal cord tissue was cut out into a block of approximately 1.5 cm x 1.5 cm x 1.5 cm that was transversely sectioned with an oscillating-blade vibratome. Most precise and even tissue sectioning was achieved by using a buffer temperature of 5 – 8°C, a cutting frequency of 100 Hz, a velocity of 2.5 mm/s and amplitude of 0.4 mm. A section thickness of 300 µm has proven to be optimal for subsequent sample mounting and indentation measurements. The acute spinal cord tissue sections were incubated in aCSF on ice until further processing for indentation measurements.

### Tissue sample mounting for indentation measurements

Spinal cord sections selected for indentation measurements were immobilized on tissue culture plastic (TCP) with Histoacryl® (B. Braun, 9381104) which was sparsely applied between the TCP and the agarose embedding at a distance of about 3 mm from the spinal cord tissue. The tissue sections were submerged in cooled aCSF during indentation measurements.

### Spinal cord transection

Zebrafish were anaesthetized by immersion in Ethyl 3-aminobenzoate methanesulfonate (0.02% in PBS, pH 7.5, Sigma-Aldrich, A5040) until respiratory movements of the opercula stopped (approximately 5 min). The surgical procedure was carried out as described in [12]. The vertebral columns were cut halfway between the dorsal fins and the opercula approximately at the level of the eighth vertebra. To account for changes of mechanical properties of the spinal cord tissue due to pre- and postoperative care, anesthesia and the incisional trauma, sham animals underwent the same procedure except for the spinal cord transection. Spinal cord transected and sham-operated fish were sacrificed and subjected to the aforementioned preparation procedure at 2 weeks post-injury (wpi), 4 wpi and 6 wpi. All zebrafish used for spinal cord transections were six to nine months old.

### Atomic force microscopy setup

Indentation measurements enabled by atomic force microscopy (AFM) and simultaneous fluorescence microscopy was performed with the CellHesion200 equipped with a motorized precision stage (JPK Instruments, Berlin) and the upright Axio Zoom.V16 stereo microscope with a PlanApo Z 0.5x objective (Carl Zeiss Microscopy, Jena). For indentation experiments, polystyrene beads (d = (37.3 ± 0.3) µm, Microparticles GmbH, PS-F-37.0) were glued to tipless silicon cantilevers (Arrow-TL1, NanoWorld) with epoxyglue. Cantilevers were calibrated using the thermal noise method [30] prior to experiments; only cantilevers with spring constants between 0.015 N/m and 0.030 N/m were used.

### Indentation measurements

To obtain detailed spatial information about the mechanical properties of zebrafish spinal cord tissues, indentation measurements were carried out on transverse tissue sections obtained from distinct locations along the anterior-posterior axis of the fish. In case of uninjured specimens, tissue sections were located approximately at the level of the 4^th^, the 12^th^, the 20^th^ and 28^th^ vertebra. In case of spinal cord transected animals, tissue sections were located rostrally and caudally in proximity (< 500 µm) to the lesion site as well as distal (≈ 2.0 mm) to it. Tissue sections from sham-operated zebrafish were located at the same positions as described for spinal cord transected animals with the addition of tissue sections at the level of the lesion where the glial bridge in spinal transected fish is formed. Each tissue section was divided into nine regions of interest (ROIs) based on the fluorescence pattern of the transgenic fish line Tg(mbp:GFP, alpha1-tubulin:mls-dsRed), enabling the discrimination of white and gray matter regions. In this line, GFP is expressed under the myelin basic protein promoter and dsRed is expressed under the alpha1-tubulin promoter and coupled to a mitochondrial leader sequence enabling the distinction between white (GFP-positive) and gray (dsRed-positive) matter regions, respectively. For each ROI, a grid of indentation points covering the entire ROI was set and force-distance curves were recorded using the AFM acquisition software (JPK Instruments). The indentation force was 4 nN and the indentation speed was 10 µm/s. The number of points per grid was chosen after estimating the approximate contact area of indenter and sample to avoid overlapping contact areas of neighboring indentation spots. For ROIs 2, 4 and 5, where a squared grid was often not suitable to probe the entire region, additional indentation curves were obtained manually. The order in which individual ROIs were probed was randomized for all tissue samples probed. To discern whether zebrafish spinal cord tissue elasticity was affected by the presence of fluorophores, all experiments were complemented by indentation measurements using wild type fish. High intensity transmitted light was sufficient to recognize areas that scattered light more strongly and therefore appeared darker (white matter) and areas that appeared lighter (gray matter). All indentation measurements took place at 18°C room temperature.

### Indentation data analysis

Force-distance curves were analyzed using the JPK data processing software (JPK Instruments, Berlin, Germany) in which the indentation segments of the approach curves are fitted with the Hertz model for a spherical indenter:

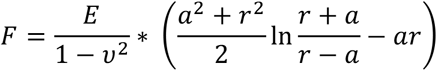

with

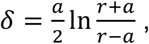

where *F* denotes the indentation force, *δ* the indentation depth, *r* the indenter radius and *a* the radius of the circular contact area between indenter and sample [31–33]. The Poisson’s ratio *ν* was set to 0.5 for all analyses. The Young’s modulus, or elasticity, *E* was used as the fitting parameter and served as a measure for the apparent elastic resistance of the probed sample to deformation. Since both measurement and analysis described in this study approximate the tissue as a purely elastic solid and do not account for viscous material properties, the values of the Young’s modulus are termed ‘apparent’. The elasticity maps were analyzed and assembled with a custom-written MATLAB algorithm (The MathWorks, Natick, MA) that used the Hertz model for a conical indenter:

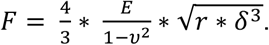

### Quantification of cell number density

Transverse tissue sections were obtained from spinal cord transected and sham-operated zebrafish (Tg(mbp:GFP, alpha1-tubulin:mls-dsRed)) at 2 wpi, 4 wpi and 6 wpi. The animals were sacrificed as described above followed by subsequent administration of PBS and 4% PFA through the *bulbus arteriosus* and incubation overnight in the same fixative. Spinal cord tissues were then dissected, post-fixed with fresh 4% PFA for 2 h and transferred to PBS. The fixed tissue samples were embedded in agarose, vibratome-sectioned and subjected to several washing and permeabilization steps using PBSTx (0.1% Triton X-100 in PBS) before nuclear staining with 4’,6-diamidino-2-phenylindole (DAPI, 1:2000 in PBS). Tissue sections were then mounted and imaged by confocal fluorescence microscopy using the Zeiss LSM700, a 20x/0.8 objective to image entire cross-sections and a 63x/1.4 objective to resolve single nuclei in individual ROIs. Fluorescence signals acquired from DsRed and GFP channels were used to identify individual gray and white matter regions as described for indentation measurements. Fluorescence signals from the DAPI channels were used to assess the number of nuclei, their volumes and cell number densities. For this purpose, acquired z-stacks of individual ROIs were converted to z-stacks of binary images by applying a Gaussian blur filter (sigma = 2), background subtraction and thresholding. A 3D objects counter [34] was then used to count nuclei detected within the imaged volume and quantify their volumes. Image processing was carried out with Fiji [35] using the same settings for all samples. Images obtained with the 63x objective were additionally subjected to a size filter to exclude background signals that could not be identified as nuclei. ROIs extracted from overview images obtained with the 20x objective yielded nuclei volumes that were summed and normalized to the volume of the imaged ROI resulting in relative nuclei densities. Volumes of single nuclei obtained with the 63x objective were used to determine an average nuclear volume for each ROI. Cell number density was then calculated by dividing the relative nuclei densities by the respective averaged volumes for single nuclei.

### Quantification of fluorescence intensity

Images from spinal cord cross-sections obtained with the 20x/0.8 objective were divided into nine ROIs as described above. The z-axis profile tool in Fiji [35] was used to identify the optical slice of the z-stack of each ROI that exhibited the highest fluorescence intensity. This slice was extracted as a 2D gray scale image and used to calculate area, mean gray values and integrated density values. Integrated density values were corrected for background fluorescence intensities. The mean fluorescence intensity values of the background were determined by obtaining images of regions not belonging to the spinal cord tissue and subjecting them to the aforementioned procedure.

### Cell Body Area Quantification

To quantify the area of individual cell bodies, confocal stacks from ROIs 1 and 4 (gray matter) and ROIs 2 (white matter) that were imaged with the 63x/1.4 objective for the cell body density assessment were reanalyzed. ROI 1, 2 and 4 were chosen as representative regions. We used the DAPI signal to detect the presence of nuclei and, hence, a cell body. The area around each nucleus devoid of GFP was outlined with the selection tool in Fiji [35], quantified and used as an indicator for cell body area.

### Graphical representation of data and statistical analysis

Each spinal cord section was divided into nine ROIs and each ROI was probed by indenting at least five different positions. For symmetry plots, the mean apparent Young’s modulus of each ROI and each animal was calculated. For gray and white matter comparisons, the mean apparent Young’s modulus for each animal was calculated from pooled gray matter ROIs (ROI 1, ROIs 3, ROIs 4) and pooled white matter ROIs (ROIs 2, ROIs 5), respectively. The sample size *n* refers to the number of animals, so that, for symmetry plots, each sample (each ROI) comprises at least 5\**n* indentations. In plots comparing combined gray and combined white matter, each sample (type of matter) contains at least 25\**n* indentations for gray matter and at least 20\**n* indentations for white matter. The graphical data representation was compiled with Python and utilizes violin plots in which dashed and full lines indicate interquartile ranges and medians, respectively. For each sample, the kernel density estimate was calculated using a bandwidth *h* with

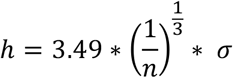

where *n* denotes the sample size and *σ* denotes the sample standard deviation [36]. For multiple comparisons, statistical analysis was performed with the Kruskal-Wallis test followed by the Dunn-Šidák post-hoc method. For pairwise comparisons, the Mann-Whitney test was used. Data obtained from confocal microscopy were processed, plotted and subjected to statistical analyses in the same manner as mechanical data. Asterisks indicate significance levels as follows: *p < 0.05; **p <0.01; ***p < 0.001, ****p < 0.0001.

## Results

### Indentation measurements on acute zebrafish spinal cord sections

To assess the mechanical properties of spinal cord tissues from adult zebrafish, we performed AFM-based indentation tests on acute, living (non-fixed), transverse spinal cord sections. These sections were obtained from vertebral levels that correspond to the 4^th^, the 12^th^, the 20^th^ and the 28^th^ vertebra (Fig.1a, Suppl.Fig.1). Each tissue section was divided into nine regions of interest (ROIs) based on the fluorescence pattern (Fig.1b) of the transgenic fish line Tg(mbp:GFP, alpha1-tubulin:mls-dsRed). The ROIs corresponded furthermore to distinct structural features of the spinal cytoarchitecture as described in [37]. ROI 1 comprises the dorsal horn (dh, Fig.1b), ROI 2 contains the dorsal longitudinal fasciculi (dlf, Fig.1b), ROI 3 and 4 comprise the ventral horns (vh, Fig.1b) and ROI 5 contains the ventral (vlf, Fig.1b) and medial longitudinal fasciculi (mlf, Fig.1b).

**Figure 1.**
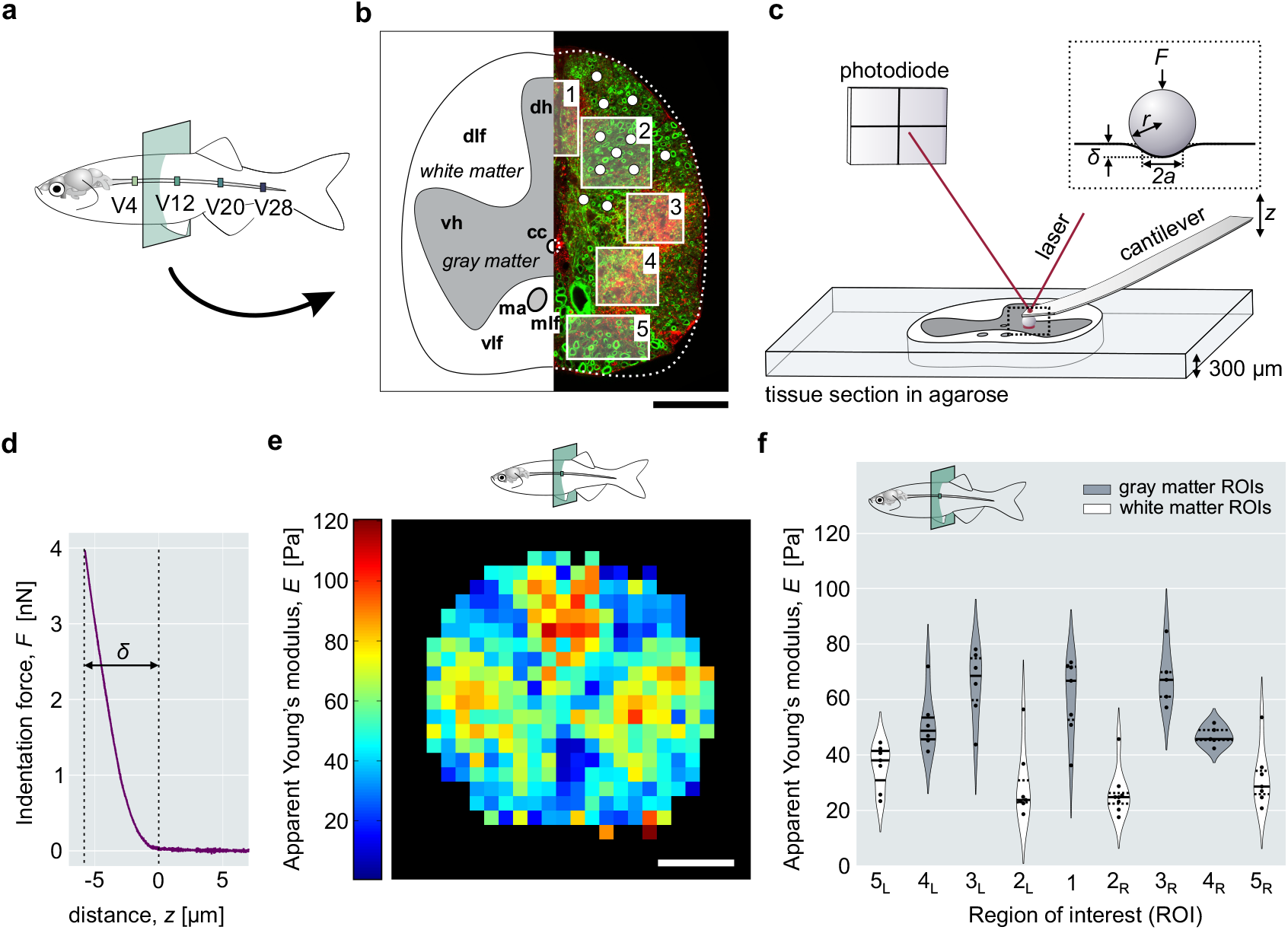
AFM-based indentation measurements on acute zebrafish spinal cord sections. **a**) Schematic representation of the adult zebrafish spinal cord with four vertebral levels from where investigated tissue sections were obtained. **b**) Schematic and fluorescence image of a transverse spinal cord section depicting gray and white matter with distinct anatomical structures and corresponding regions of interest. Exemplary indentation spots (white) are shown for ROI 2. Myelin-rich regions are labeled with GFP (green, white matter). Mitochondria-rich regions are labeled with dsRed (red, gray matter). (dh: dorsal horn, vh: ventral horn, dlf: dorsal longitudinal fascicle, vlf: ventral longitudinal fascicle, mlf: medial longitudinal fascicle, ma: Mauthner axons, cc: central canal). **c**) Schematic of an AFM-based indentation setup. The insert shows the indentation force, *F*, and the geometrical factors used by the Hertz model: indentation depth *δ*, radius of the indenter *r* and radius of the contact area *a*. **d**) Exemplary force-distance curve with indentation depth, *δ*. **e**) Elasticity map showing the spatial distribution of apparent Young’s moduli of an entire spinal cord cross-section obtained from the level of the 12^th^ vertebra. **f**) Violin plot showing the apparent Young’s moduli of individual ROIs from one tissue section obtained from the level of the 12^th^ vertebra. Here, each data point represents the apparent Young’s modulus from one indentation spot. Enumeration of individual ROIs was carried out as shown in b). Indices L and R denote the left and right sides of the tissue section, respectively. Scale bars, 100 µm.

The tissue sections were probed (Fig.1c) using a spherical indenter. Each indentation yielded a force-distance curve (Fig.1d) that was fitted using the Hertz model (Fig.1c, insert) to derive the apparent Young’s modulus *E* [31, 32] (see Methods). Elasticity maps (Fig.1e) of the entire spinal cord cross section were recorded to display the distribution of apparent Young’s moduli in a color-coded manner. To compare stiffness values of different ROIs and compare their distribution, apparent Young’s moduli were plotted as violin plots (Fig.1f). Both modes of data presentation suggest that gray matter regions are stiffer than white matter regions in the displayed sample section, which we set out to further investigate.

As the mechanical characterization of zebrafish spinal cord tissue reported here aimed at describing intrinsic mechanical tissue properties that occur in living tissues, viability and structural integrity of the tissue sections were investigated with necrosis and apoptosis assessments (see Suppl. Methods and Suppl. Information). The results obtained from both viability assays showed that the sample preparation and course of the experiment did not impair the overall tissue viability or tissue architecture in the time interval of 5 hours *post-mortem* used for mechanical testing (Suppl.Fig.1).

### Gray matter regions are stiffer than white matter regions

To systematically characterize the mechanical phenotype of spinal cords in adult zebrafish, we performed indentation measurements on individual tissue sections of ten zebrafish (Fig. 2a) that were obtained from different vertebral levels (Fig. 2b) and repeated those experiments for zebrafish of different ages (Fig. 2c). Regions corresponding to gray matter (ROIs 1, 3, 4) were stiffer than regions corresponding to white matter (ROIs 2, 5) (Fig. 2a). Within the gray matter, the dorsal horns (ROIs 1) and the most dorsal parts of the ventral horns (ROIs 3) displayed greater median elasticities in comparison to ROIs 4. White matter regions ROIs 2 showed smaller median elasticities than white matter regions ROIs 5. The apparent Young’s moduli of all individual ROIs were distributed symmetrically with respect to the dorso-ventral midline. This bilateral symmetry reflected the tissue’s architectural symmetry as indicated by the observed fluorescence pattern (Fig. 1b). The mechanical symmetry was maintained in all investigated tissue sections obtained from uninjured animals along the anterior-posterior axis of the spinal cord (Suppl. Fig. 3,4) regardless of age or presence of fluorophores (Suppl. Fig. 3,4,5).

**Figure 2.**
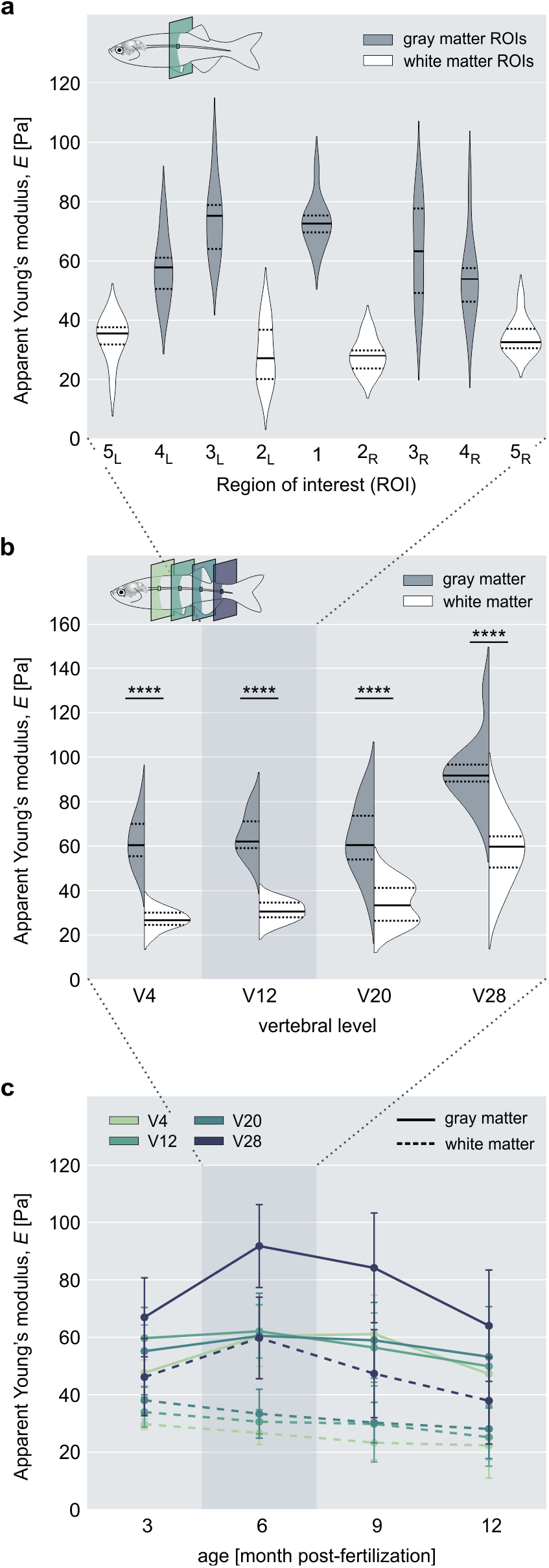
Spinal cord elasticity of adult zebrafish. **a**) The apparent Young’s moduli of individual ROIs from spinal cord sections of ten animals (*n* = 10) located at the level of the 12^th^ vertebra. **b**) Distribution of apparent Young’s moduli of combined gray and white matter ROIs along the anterior-posterior axis at the indicated vertebral levels. All specimens (*n* = 10) were 6 months old. Solid lines and dashed lines represent medians and interquartile ranges, respectively. ****p < 0.0001. **c**) Comparison of the apparent Young’s moduli of spinal gray and white matter at different vertebral levels (V4 - V28) and at 3 months post-fertilization (mpf) (*n* = 5), 6 mpf (*n* = 10), 9 mpf (*n* = 10) and 12 mpf (*n* = 10).

### The difference between gray and white matter is maintained along the A-P axis

To assess the distribution of elasticity values at different vertebral levels, we combined ROIs 1, 3 and 4 to gray matter and ROIs 2 and 5 to white matter and plotted their respective elasticity values as a function of location along the anterior-posterior axis of the spinal cord (Fig. 2b). Spinal gray and white matter within the 4^th^, the 12^th^, the 20^th^ vertebra displayed comparable elasticity values respectively, whereas both gray and white matter showed elevated values near the 28^th^ vertebra (Fig. 2b,c, Suppl. Fig. 3,4).

The difference of apparent Young’s moduli between gray and white matter remained constant along the anterior-posterior axis (Fig. 2b) and was furthermore maintained throughout their life span (Fig. 2c), although absolute elasticity values of both gray and white matter decreased with age (Fig. 2c).

To test whether the presence of transgenic fluorophores might have an influence on this pattern, we also measured wild type fish. In those specimens, the absolute apparent Young’s moduli of gray matter, but not white matter were lower as compared to transgenic fish (Suppl. Fig. 5a-d). Wild type spinal tissue also showed a less pronounced increase of elasticity and difference between white and gray matter in tissue sections located at the 28^th^ vertebra (Suppl. Fig. 5e) and the difference between spinal gray matter and spinal white matter was also maintained throughout their life span (Suppl. Fig. 5f).

### Adult zebrafish spinal cord regeneration is accompanied by transient tissue stiffening

To capture a spatially and temporally resolved mechanical phenotype in response to spinal cord injury, indentation measurements were executed with tissue sections from different spinal levels (see Methods and Suppl. Fig. 1) at 2 weeks post-injury (wpi), 4 wpi and 6 wpi. In spinal transected fish, the vertebral column was severed halfway between the dorsal fins and the opercula, which corresponds to the location of the 8^th^ vertebra [12]. The apparent Young’s moduli of gray and white matter from uninjured zebrafish displayed constant elasticity values around this spinal level and were used as a reference for both spinal cord transected and sham-operated fish (Suppl. Fig. 2,3). After complete spinal cord transection, caudal spinal cord sections proximal to the lesion site displayed a significant increase of white matter stiffness at 2 wpi which rendered the formerly pronounced mechanical difference between gray and white matter non-significant (Suppl. Fig. 6f). By 4 wpi, the apparent Young’s moduli of both gray and white matter had increased further in comparison to values measured at 2 wpi and to values from uninjured control animals (Suppl. Fig. 6f). White and gray matter reached comparable elasticity levels (Fig. 3a, Suppl. Fig. 7e). At 6 wpi, the elasticity of both gray and white matter declined as compared to 4 wpi, but remained elevated with respect to uninjured control animals (Suppl. Fig. 6f). The difference between gray and white matter elasticity was re-established (Suppl. Fig. 6f, 7f). Rostral tissue sections that were located proximal to the lesion site displayed a similar stiffness evolution in the same time interval post-injury, albeit the changes of apparent Young’s moduli were less pronounced (Suppl. Fig. 6c). At 2 wpi, the apparent Young’s modulus of white matter increased and the mechanical difference between gray and white matter became less significant (Suppl. Fig. 6c, Suppl. Fig. 7a). At 4 wpi, the apparent Young’s moduli of gray and whiter matter increased, but gray matter remained stiffer than white matter (Fig. 3b, Suppl. Fig. 6c, Suppl. Fig. 7b). At 6 wpi, the gray and white matter elasticity decreased towards the level of uninjured animals and their characteristic difference as observed in homeostatic tissues was re-established. White matter elasticity values remained elevated in comparison to uninjured control fish (Suppl. Fig. 6c, Suppl. Fig. 7c). Spinal cord tissue sections that were located rostrally and distal to the lesion site displayed no change of mechanical properties in comparison to uninjured control animals (Suppl. Fig. 6a). Caudally, however, distal tissue sections showed an increase of white matter, but not gray matter elasticity that remained constant during the investigated time interval (Suppl. Fig. 6h). Spinal cord tissues of sham-operated zebrafish did not differ mechanically from uninjured counterpart at all investigated time points (Suppl. Fig. 6b,d,e,g,I, Suppl. Fig. 8). While absolute values, spread of data and significance levels differed in some tissue sections in wild type fish, spinal cord transections (Suppl. Fig. 9a,c,f,h, Suppl. Fig. 10) and sham treatments (Suppl. Fig. 9b,d,e,g,I, Suppl. Fig. 11) elicited a spatio-temporal profile of tissue elasticity that was comparable to their respective transgenic counterparts for all investigated locations and time points.

**Figure 3.**
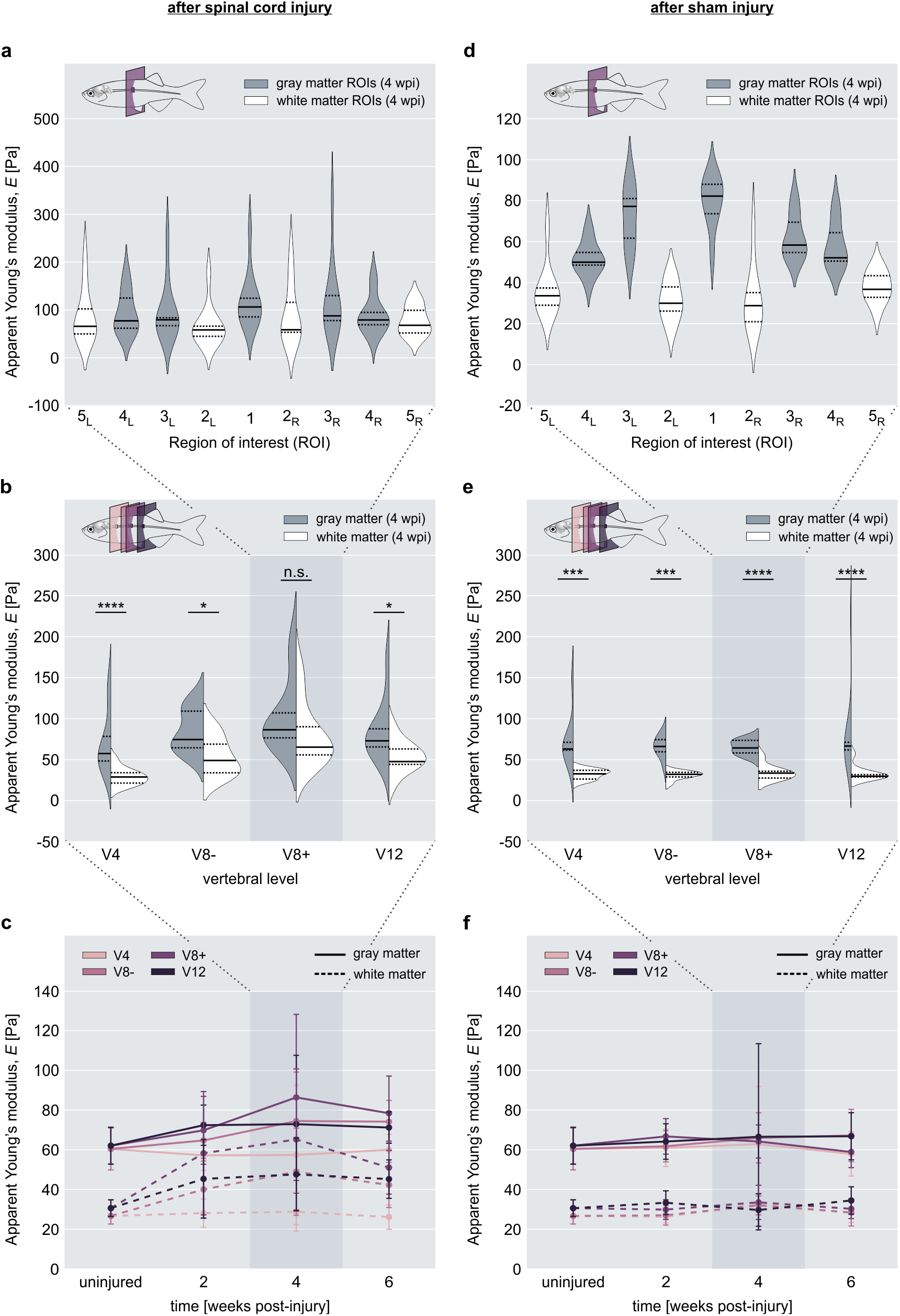
Spinal cord elasticity of adult zebrafish during regeneration. **a**) The apparent Young’s moduli of individual ROIs from spinal cord sections (*n* = 10) located caudally near the lesion site (V8+) at 4 weeks post-injury (wpi). **b**) Distribution of apparent Young’s moduli of combined gray and white matter at 4 wpi (*n* = 10). Tissue sections were located rostrally and caudally proximal (V8-, V8+) and distal (V4, V12) to the lesion site. **c**) Comparison of the apparent Young’s moduli of spinal gray and white matter from the vertebral levels indicated in b) at 2 wpi (*n* = 10), 4 wpi (*n* = 10) and 6 wpi (*n* = 10). **d**) The apparent Young’s moduli of individual ROIs from spinal cord sections (n = 10) of sham injured zebrafish at 4 wpi. The location of tissue section corresponded to the V8+ level in spinal transected animals. **e**) Distribution of apparent Young’s moduli of combined gray and white matter at 4 wpi (*n* = 9). Tissue sections were located at the same vertebral levels as tissue sections from spinal transected fish (V4, V8-, V8+, V12). **f**) Comparison of the apparent Young’s moduli of spinal gray and white matter from the vertebral levels indicated in e) at 2 wpi (*n* = 10), 4 wpi (*n* = 9) and 6 wpi (*n* = 10). Solid lines and dashed lines in a), b), d) and e) represent medians and interquartile ranges, respectively. *p < 0.05, ***p < 0.001, ****p < 0.0001.

### Cell body density inconsistently contributes to spinal cord elasticity

To explain the mechanical difference between gray and white matter areas in murine spinal cord tissues, Koser et al. have proposed to correlate the distribution and sizes of cell nuclei, as a proxy for cell number density, with the calculated apparent Young’s moduli of the respective tissue regions [38]. To be able to investigate the role of cell number density as a potential determinant of spinal cord tissue elasticity in adult zebrafish, we obtained confocal stacks of DAPI-stained tissue sections (Fig. 4a) and quantified the distribution and densities of cell bodies in uninjured spinal cord tissue and after complete transection. The use of the same transgenic fish line characterized by mechanical testing allowed the correlation between mechanical properties and cell body density in individual ROIs.

**Figure 4.**
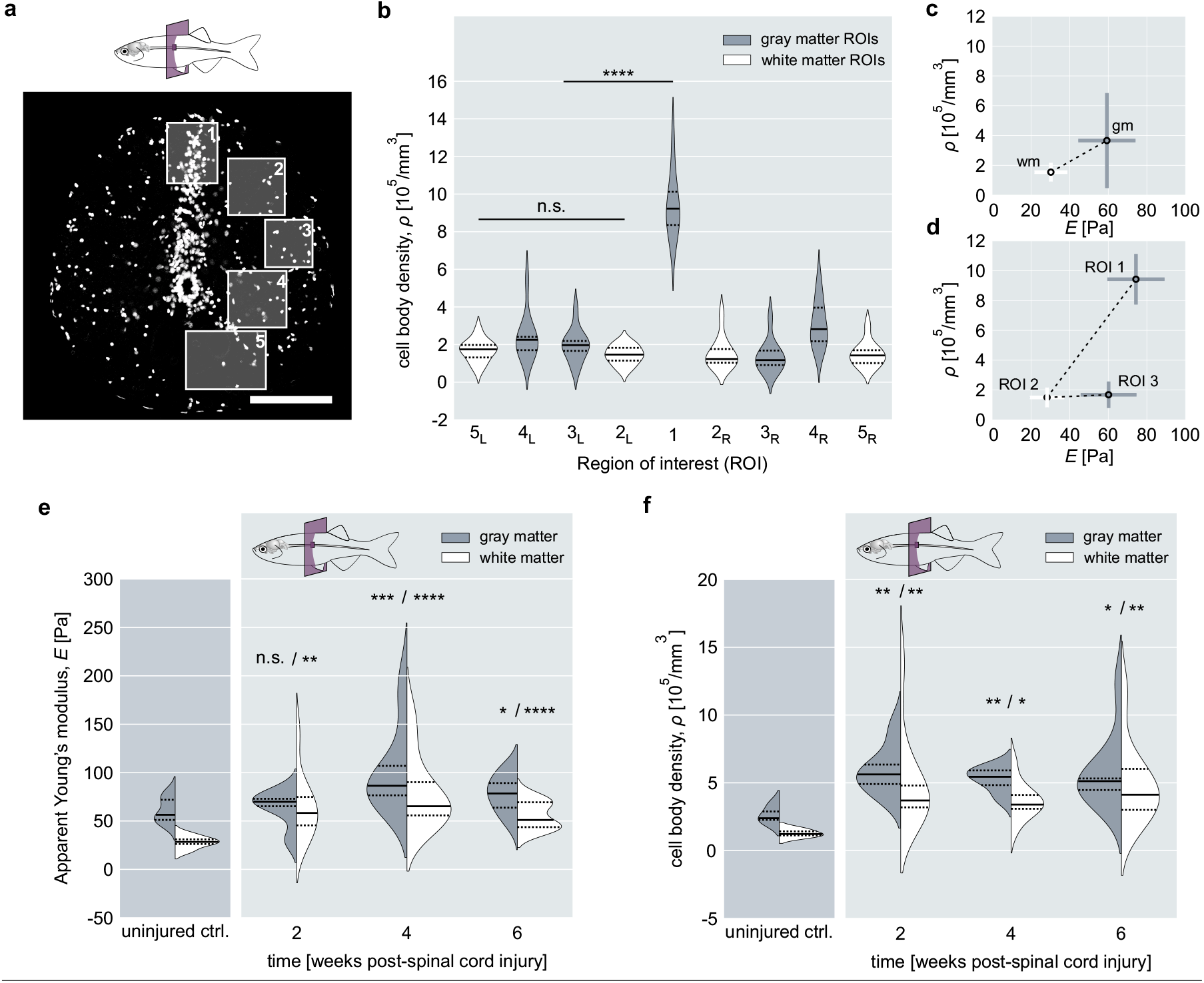
Quantification of cell body density. **a**) Fluorescence image showing the distribution of DAPI-stained nuclei (white) in a transverse zebrafish spinal cord section obtained from a level corresponding to the 8^th^-9^th^ vertebra. Scale bar, 100 µm. **b**) Distribution of cell body density in individual ROIs (*n* = 10) as indicated in a). Solid lines and dashed lines represent medians and interquartile ranges, respectively. **c**) Correlation of median apparent Young’s moduli from combined gray (gm) and combined white matter (wm) with respective cell body densities. **d**) Correlation between apparent Young’s moduli and cell body densities of ROI 1, 2 and 3. **e**) Apparent Young’s moduli of spinal gray matter and spinal white matter at the level of the 8^th^-9^th^ vertebra during regeneration at 2 wpi (*n* = 10), 4 wpi (*n* = 10) and 6 wpi (*n* = 10). Solid lines and dashed lines represent medians and interquartile ranges, respectively. Significance levels correspond to pairwise comparisons with gray and white matter from uninjured control animals (*n* = 10), respectively. **f**) Cell number density of spinal gray and white matter during regeneration at 2 wpi (*n* = 10), 4 wpi (*n* = 10) and 6 wpi (*n* = 9). Solid lines and dashed lines represent medians and interquartile ranges, respectively. Significance levels correspond to pairwise comparisons with gray and white matter from uninjured control animals (*n* = 3), respectively. *p < 0.05, **p < 0.01, ***p < 0.001, ****p < 0.0001.

In uninjured tissues, gray and white matter elasticity correlated with cell number density only when all individual gray matter and white matter regions were combined (Fig. 4b,c). The positive correlation between cell number density and the apparent Young’s moduli originated from the high density of cells in the dorsal horn (ROI 1, Fig. 4a,b). The ventral horns (ROIs 3 and 4, Fig. 4a,b) displayed an amount of cell bodies comparable to white matter regions (ROIs 2 and 5), but differed significantly in elasticity (Fig. 4d). After spinal cord injury, we observed an increase of both apparent Young’s moduli and cell body density (Fig. 4e,f). However, the exact temporal profile of cell body density did not mirror the evolution of mechanical properties in the course of regeneration. For instance, the increase in elasticity after spinal cord injury peaks at 4 wpi (Fig. 4f) for both gray and white matter, but the respective cell body densities showed a significant increase at 2 wpi that remained until 4 wpi. During regeneration, the cell body density of white matter was lowest at 4 wpi – a time point when elasticity values were highest.

### Tissue architecture correlates with mechanical differences

Confocal fluorescence microscopy using the Tg(mbp:GFP, alpha1-tubulin:mls-dsRed) fish line revealed further structural elements that might contribute to zebrafish spinal cord elasticity. In this fish line, oligodendrocytes express GFP under the myelin basic protein promoter [27], which allows the visualization of oligodendrocytes and axonal orientation in each ROI. As oligodendrocytes myelinate multiple axons, they might serve as a crosslinking factor in the spinal parenchyma. Axonal orientation has been shown to influence the tissue’s resistance to indentation as axonal projections in the white matter are aligned in parallel to the indentation direction in transverse tissue sections [38]. As exemplarily indicated for ROIs 1, 4 and 2, gray and white matter regions displayed different mechanical properties that could not be explained by their respective cell number densities alone. ROI 1 showed the highest amount of cell bodies (Fig. 4c, Fig. 5a) and the least amount of oligodendrocytes (Fig. 5a,d). ROI 4 displayed a median apparent Young’s modulus twice as high as that of ROI 2, although both regions had a similar number of cell bodies (Fig. 4c). They differed, however, with regard to axonal orientation and cell body size (Fig. 5b,e). Furthermore, we observed blood vessels predominantly in gray matter regions, but not white matter regions (Fig. 5f).

**Figure 5.**
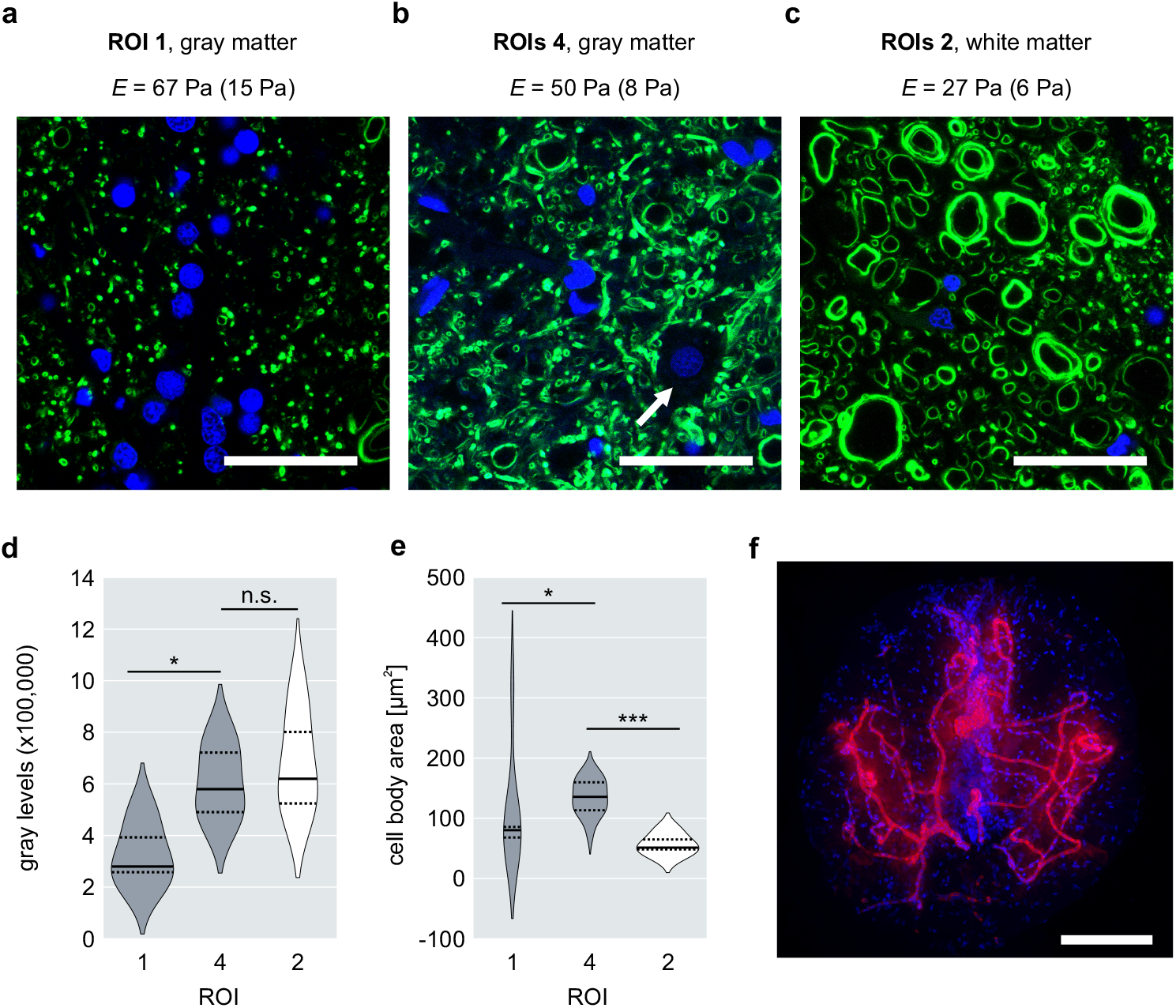
Additional contributing factors to adult zebrafish spinal cord elasticity. Exemplary fluorescence images of two gray matter regions, ROI 1 (**a**) and ROI 4 (**b**), and one white matter region, ROI 2 (**c**), showing oligodendrocytes (GFP, green) and cell nuclei (DAPI, blue). Scale bar, 50 µm. The apparent Young’s moduli for the respective ROIs are given as median (inter quartile range). **d**) Fluorescence intensity quantification of the GFP signal from ROIs 1,4 and 2 (*n* = 5). **e**) Quantification of the cell body areas in ROIs 1,4 and 2 (*n* = 7). The position and size of individual cell body areas were determined by using the DAPI signal to detect the presence of a cell body and outlining the area around each nucleus devoid of GFP (arrow in b). *p < 0.05, ***p < 0.001. **f**) Fluorescence image of a transverse spinal cord section showing the vasculature (autofluorescence, red) in the spinal gray matter of adult zebrafish. Cell nuclei are depicted in blue. Scale bar, 100 µm.

## Discussion

The present study focused on the mechanical characterization of the adult zebrafish spinal cord in order to investigate the evolution of the mechanical tissue properties following spinal cord transection. An efficient and reliable protocol was established that allowed obtaining and preserving acute zebrafish spinal cord sections for consecutive mechanical measurements without fixation. Since the sectioned spinal cord tissues remained viable and structurally intact during the time interval of mechanical testing, we consider our results to be physiologically relevant. AFM based indentation measurements were then employed to determine the apparent Young’s moduli of spinal cord tissues from uninjured zebrafish and at distinct time points during regeneration. We showed that the apparent Young’s moduli of gray matter regions were greater than those of white matter regions in spinal cords of uninjured, adult zebrafish (Fig. 6a). Previous studies measuring the stiffness of nervous tissues from mouse and rat also found that gray matter was stiffer than white matter [24, 38, 39]. However, the absolute values of zebrafish spinal cord tissue stiffness were almost an order of magnitude lower as compared to their rodent counterparts [24, 38]. The distribution of individual regions of interest furthermore mirrored the symmetry of the spinal cytoarchitecture of adult zebrafish as described in [37]. Each tissue region contains a specific set of cell bodies, processes and filaments that vary in structural arrangement and/or density and consequently amount to distinct tissue architectures. Gray matter regions consist of densely packed processes, synapses, neurofilaments and neuronal cell bodies, whereas white matter regions contain ascending and descending myelinated axon tracts that run in parallel to the anterior-posterior axis of the spinal cord and are crosslinked by oligodendrocytes [37]. The dorsal horn exhibited the highest elasticity values in uninjured specimen and displayed a tissue architecture that was dominated by the presence of densely packed cell bodies. In addition, blood vessels are present predominantly in gray matter regions of the zebrafish spinal cord where they might add structural support and contribute to the greater tissue elasticity values measured.

**Figure 6.**
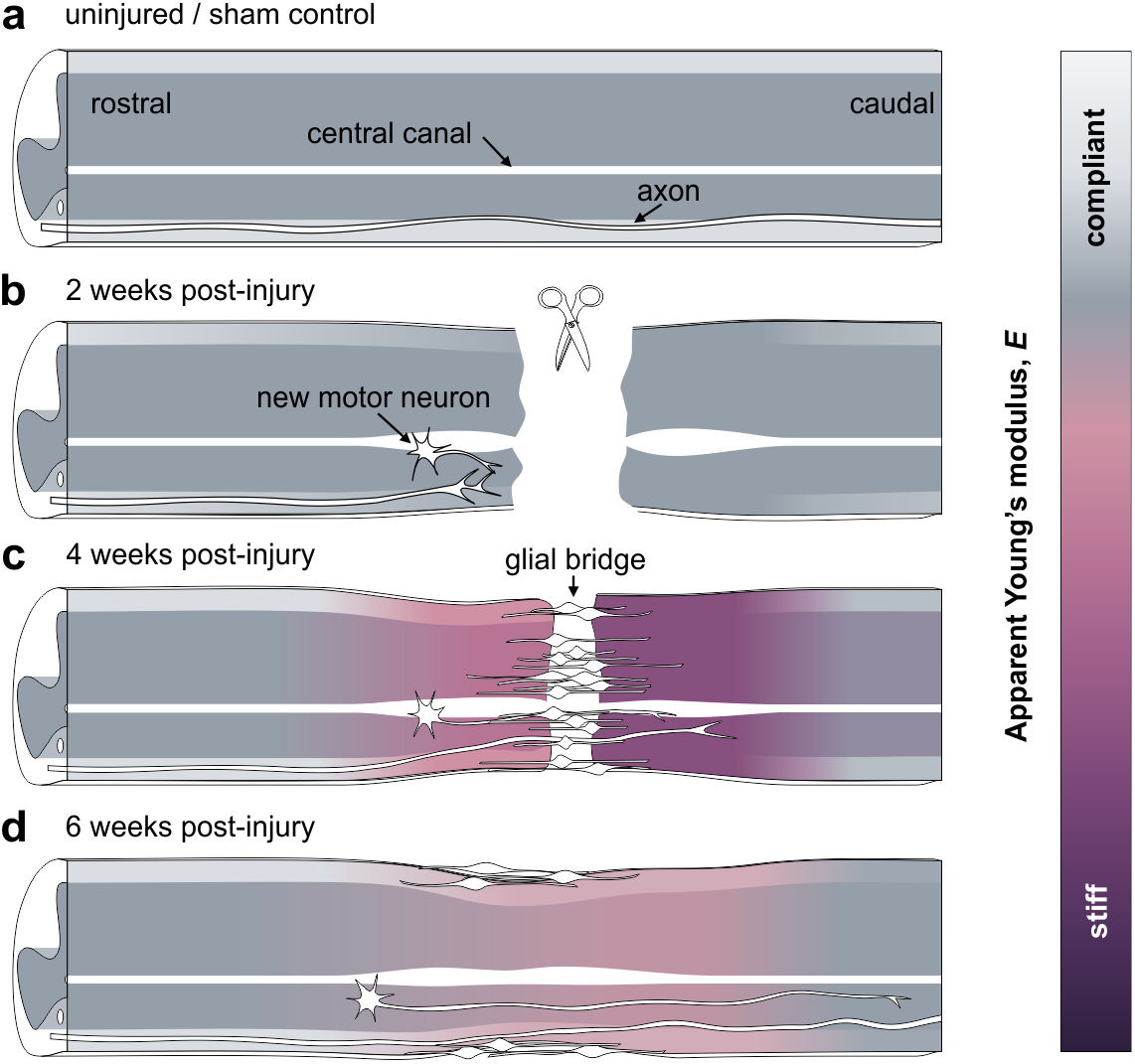
Graphical summary of the spatio-temporally changing mechanical phenotype of adult zebrafish during spinal cord regeneration. a) Schematic representation of an uninjured or sham operated spinal cord tissue in which gray matter was stiffer than white matter along the anterior-posterior axis. b) At 2 wpi, white matter regions in proximity to the lesion site had increased in stiffness. Gray matter remained mechanically at uninjured levels. c) At 4 wpi, both white and gray matter stiffened significantly and, in caudal tissue sections, became mechanically comparable. This effect was more pronounced in proximity to the lesion site than in distal tissue sections. c) At 6 wpi, the elasticities of gray and white matter had decreased in comparison to 4 wpi, but remained elevated in comparison to uninjured control animals. The mechanical difference between gray and white matter was re-established.

The region located circumferentially around the central canal was subjected to indentation measurements as well, but the vast majority of force-indentation curves recorded in this region could not be analyzed due to missing baselines (approach segments) or multiple slopes of indention segments. This effect could be explained by uneven sectioning results that were likely caused by the mechanical instability of columnar epithelium lining the central canal. This particular region was therefore excluded from data presentation and discussion.

Median elasticities of gray and white matter regions were furthermore comparable in most vertebral levels possibly reflecting the likewise comparable tissue architecture in the spinal cord along the length of the animal. The increased apparent Young’s moduli of tissue sections located at the 28^th^ vertebra might arise from a more densely packed spinal cord tissue due to the caudally decreasing diameter of the spinal cord and/or altered structural properties of individual tissue components. For instance, sensory fibers and processes of radial glia, both of which display progressively increasing densities toward the caudal part of the spinal cord [37], might serve to explain the increase of tissue elasticity in ROIs 5. Apart from that, it is currently not known if and how the spinal composition changes in the most caudal spinal cord parts.

After complete spinal cord transection, adult zebrafish displayed an increase of spinal gray matter and spinal white matter elasticity that transiently obliterated both the distinct mechanical difference between the two types of matter and, in caudal sections, the mechanical symmetry with respect to the dorso-ventral midline. This effect was more pronounced in the vicinity of the lesion site and less marked in tissue sections located distal to the lesion site (Fig. 6b-d). As all sham treatments, i.e. pre- and postoperative care, anesthesia and the incisional trauma, could be excluded as potential elicitors, the change of mechanical properties in the course of regeneration can be attributed to the spinal cord transection. This suggests a correlation between the cellular events induced by the lesion and the spatio-temporal profile of tissue elasticity in the course of regeneration. In light of recent studies on neural mechanosensitivity and stiffness changes of nervous tissue during developmental and pathological events, it is an open question if and how mechanical cues are provided by the spinal cord tissue after spinal cord injury. It has been speculated that the glial-fibrotic scar in mammals — which is densely packed with cells and ECM — is significantly stiffer than its environment and may act as a mechanical and structural barrier obstructing the penetration of axons and thereby axonal regrowth across the lesion site [2, 40]. In contrast, adult zebrafish show robust functional regeneration and tissue repair after spinal cord injury [6], and the apparent Young’s moduli of both gray and white matter regions reached their highest level at a time point when regrowing axons traverse the injury site. Koser et al. have shown that growing axons of retinal ganglion cells possess mechanosensitive ion channels that allow them to detect differences in tissue stiffness in the developing frog brain and, as a result, extend faster, straighter and in a fasciculated way [18]. A similar mechanism might be at play in the regenerating zebrafish spinal cord in which the exposure of regrowing axons to greater tissue stiffness may promote axonal pathfinding by enhanced fasciculation and growth velocity. In any case, the fact that spinal cord regeneration in zebrafish was accompanied by an increase in tissue elasticity motivates to question the assumption that axonal regrowth after spinal cord injury in mammals is impeded by the presumably stiffer environment of the glial-fibrotic scar [2]. In fact, Moeendarbary et al. have recently shown that neural tissue in rat brain cortex and spinal cord softens after traumatic injury, which correlates with an increase of expression levels of glial intermediate filaments and ECM components [2, 24]. Their finding and ours combined supports the hypothesis that an increase rather than a decrease in spinal cord elasticity after injury might facilitate neuronal regrowth and spinal cord regeneration.

An additional indicator for mechanically guided axonal pathfinding in regenerating zebrafish might be presented by axonal rerouting. As described in [41], regrowing axons that descend from the brainstem encounter remaining myelin debris and reroute from the white matter to the central gray matter caudal to the lesion site. The observed rerouting lets the authors suggest that degenerating tracts are not a preferred substrate for axonal regrowth, although it has been shown that myelin-associated inhibitors of regeneration in mammals are growth-permissive in zebrafish [42, 43]. In light of the presented mechanical characterization, it seems possible that the observed rerouting is a consequence of altered mechanical tissue properties and axonal mechanosensing.

While the mechanical properties of uninjured, spinal transected and sham operated zebrafish were qualitatively comparable between transgenic and wild type fish for the majority of investigated locations and time points, absolute values, spread of data and significance levels differed in some tissue sections. This might be in part attributable to the lower number of investigated wild type specimens and the concomitant impact of intraspecies variation on the small-sized population. However, it cannot be ruled out that the presence of fluorophores influences the mechanical properties of the spinal cord tissue and therefore leads to differing absolute values of apparent Young’s moduli.

The mechanical properties of biological tissues are thought to be determined by the material properties of the constituent cells, ECM and the degree of intercellular adhesion and connectivity [19]. Koser et al. have proposed to consider additional criteria to predict nervous tissue stiffness based on fluorescently labeled tissue components such as cell body density, myelin content, collagen content, extra cellular matrix composition and axonal orientation [20, 38]. Our results show that cell body density is not sufficient to explain the mechanical differences between gray and white matter in zebrafish spinal cords nor the spatio-temporal evolution of the apparent Young’s modulus during spinal cord regeneration. Based on our microscopy analyses and previously published reports [8, 37, 41, 44], we submit that spinal cord tissue elasticity in zebrafish results from a complex synergistic effect in which cell body density and concomitant packing density, single cell stiffness, degree of crosslinking, vasculature and the presence and composition of ECM and/or yet undetermined factors contribute differently at distinct time points post-injury. Additional information about the structural determinants of zebrafish spinal tissue elasticity might be obtained from indentation measurements using tissues that were sectioned along sagittal or coronal planes. As described in [38], murine spinal white matter regions behave transversely isotropic which has been attributed to the different orientation of axon bundles as they are aligned perpendicular to the indentation direction in coronal and sagittal sections, but parallel to it in transverse sections. Indentations executed on transversely sectioned white matter regions might elicit a buckling response instead of or in addition to compression and would therefore yield underestimated elasticity values. This might contribute to the lower apparent Young’s moduli of white matter regions in both mice and zebrafish spinal cords. However, the zebrafish spinal cord is significantly smaller in diameter than its murine counterpart, which aggravates precise vibratome-sectioning along coronal or sagittal planes. Lipid-rich membranes might also cause the comparatively low elasticity values of white matter regions. These surround myelinated axons and close in dome-like structures upon sectioning. Such structures might yield a slippage of the indenter and prevent proper indentation if the indentation force and indentation depth are too small (Suppl. Fig. 12). Therefore, future experiments may involve non-invasive techniques that obviate the need for tissue sectioning or even allow measurements along different anatomical planes *in vivo*. Non-invasive methods such as confocal Brillouin microscopy and magnetic resonance elastography have been applied to many complex materials in order to extract physical properties such as tensile and compressive strain, elastic moduli and viscosity [22, 45–47]. Further technological advancements might render these methods applicable to adult zebrafish and expand our current understanding of mechanical signaling during spinal cord regeneration.

Moreover, AFM-based indentation experiments require a rather invasive sample preparation that includes the isolation of the spinal tissue from the organism and subsequent sectioning. While we have shown that our preparation preserves the viability of the cells in the tissue, these preparatory procedures could already alter the mechanical properties of the spinal cord present *in vivo*. For instance, the zebrafish spinal cord as a whole may be under tension in the living organism, which would not be detectable after dissection or tissue sectioning. The same argument is true for pulsatile blood flow through vessels that penetrate the spinal parenchyma. These factors contribute to the mechanical tissue properties, might provide additional signals to maintain cellular and tissue homeostasis, and might likewise affect tissue regeneration and repair. Gefen and Margulies, for instance, showed that the mechanical properties of brain tissue *in vivo* differ from excised tissue only after repetitive, but not at the first indentation [48]. Weickenmeier et al. used magnetic resonance elastography and showed that brain tissue rapidly stiffens after death [49]. As it is currently unclear how *in vivo* mechanical properties change post mortem, mechanical tissue properties must be measured *in vivo* to rule out any effects elicited by the death of the animal. In fact, a recent publication from our lab employed a custom-built confocal Brillouin microscopy setup and showed that spinal cord injury and repair in living zebrafish larvae coincides with significant Brillouin shift changes. After spinal cord transection, the Brillouin shifts, corresponding to the longitudinal elastic modulus of the material, measured in the lesion site decreased and gradually increased thereafter [47]. This seemingly contrasts our AFM-based indentation results obtained *ex vivo.* However, both studies differ with respect to multiple experimental parameters such as the age of the specimens and possibly corresponding regenerative capacities or direction of measurement. Furthermore, zebrafish must be sedated by immersion in tricaine containing media for confocal Brillouin microscopy, which could induce acidification and change mechanical tissue properties as previously reported [50, 51]. Yet another aspect is the interpretation of the longitudinal modulus acquired by Brillouin microscopy as it measures on very short time (GHz) and length scales, whereas AFM provides information on much larger time (Hz) and length scales. Thus, longitudinal and Young’s modulus must not necessarily correlate (for further discussion see [47]). However, as mechanical measurements appear to be dependent on several experimental parameters, the above-mentioned controversies emphasize the need for further investigations of mechanical tissue properties across all spatio-temporal scales.

## Conclusion

Traumatic spinal cord injury in humans is accompanied by an irreversible impair of information exchange between the brain and the periphery of the body. Depending on the severity of the injury and its location on the spinal cord, patients may suffer from a partial or complete loss of sensory function and motor control of extremities and/or body compartments. While great effort has been made to identify and eradicate biochemical signals that are inhibitory for nerve fiber growth, functional repair and tissue restoration, there is no treatment yet to achieve complete recovery after spinal cord injury [52]. The mechanical characterization of the regenerating, adult zebrafish spinal cord, as presented in this study, constitutes a novel, interdisciplinary approach to assess and interpret the mechanisms that govern the cellular response during successful spinal cord repair. It is the first detailed study of the evolution of mechanical properties in adult zebrafish after spinal cord injury and documents the spatio-temporal changes of mechanical tissue properties provided by the spinal cord tissue to nerve cells and supporting cell residing in the tissue. This work will serve as a basis for future studies to link spinal cytoarchitecture before and after injury to tissue stiffness with the ultimate goal of tuning spinal cord tissue mechanics towards successful functional repair also in humans.

## Author contributions

J.G., S.M., M.B. and V.K. conceived the project. S.M. designed and performed sample preparation, indentation experiments, and data evaluation. S.M. and M.K. performed immunohistochemistry and imaging. M.K. performed image analyses. S.M. and T.H. performed viability assays. S.M. performed animal experiments. S.A. developed a python script for data representation. A.T. and V.K. advised on animal experiments and indentation measurements, respectively. M.B. provided animals and methodology for animal experiments. J.G. supervised the project. S.M. and J.G. wrote the manuscript. All authors reviewed the manuscript.

## Acknowledgements

We thank the fish facility at BIOTEC, TU Dresden for excellent fish care, JPK Instruments, Berlin for technical support, the light microscopy facility at CMCB, TU Dresden, and Benoit Lombardot, MPI CBG Dresden, for support with image analysis. We thank Andreas Christ for the MatLab Script for elasticity map analysis. We thank Daniel Wehner, Michell Reimer, Kristian Franze, Andreas Christ and Elke Ulbricht for fruitful discussions. Work by V.K. and M.B. was supported by the European Union (ERA-NET NEURON NeuroNiche project 01EW1708) and ERC advanced grant (Zf-BrainReg) to MB. Financial support from the Alexander-von-Humboldt Stiftung (Humboldt-Professorship to J.G.) and the Sächsisches Ministerium für Wissenschaft und Kunst (European Fund for Regional Development — EFRE to the light microscopy and electron microscopy facilities at CMCB, TU Dresden) is gratefully acknowledged.

## References

1. Verma, P. and J. Fawcett, Spinal cord regeneration. Adv Biochem Eng Biotechnol, 2005. 94: p. 43–66.

2. Fawcett, J.W. and R.A. Asher, The glial scar and central nervous system repair. Brain Res Bull, 1999. 49(6): p. 377–91.

3. Cregg, J.M., et al., Functional regeneration beyond the glial scar. Exp Neurol, 2014. 253C: p. 197–207.

4. Silver, J. and J.H. Miller, Regeneration beyond the glial scar. Nat Rev Neurosci, 2004. 5(2): p. 146–56.

5. Vajn, K., et al., Temporal profile of endogenous anatomical repair and functional recovery following spinal cord injury in adult zebrafish. PLoS One, 2014. 9(8): p. e105857.

6. Becker, C.G. and T. Becker, Model Organisms in Spinal Cord Regeneration. 2007: Wiley.

7. Hui, S.P., A. Dutta, and S. Ghosh, Cellular response after crush injury in adult zebrafish spinal cord. Dev Dyn, 2010. 239(11): p. 2962–79.

8. Reimer, M.M., et al., Motor neuron regeneration in adult zebrafish. Journal of Neuroscience, 2008. 28(34): p. 8510–8516.

9. Goldshmit, Y., et al., Fgf-Dependent Glial Cell Bridges Facilitate Spinal Cord Regeneration in Zebrafish. Journal of Neuroscience, 2012. 32(22): p. 7477–7492.

10. Becker, T. and C.G. Becker, Regenerating descending axons preferentially reroute to the gray matter in the presence of a general macrophage/microglial reaction caudal to a spinal transection in adult zebrafish. J Comp Neurol, 2001. 433(1): p. 131–47.

11. Kuscha, V., et al., Lesion-induced generation of interneuron cell types in specific dorsoventral domains in the spinal cord of adult zebrafish. Journal of Comparative Neurology, 2012. 520(16): p. 3604–3616.

12. Becker, T., et al., Axonal regrowth after spinal cord transection in adult zebrafish. Journal of Comparative Neurology, 1997. 377(4): p. 577–595.

13. Wehner, D., et al., Wnt signaling controls pro-regenerative Collagen XII in functional spinal cord regeneration in zebrafish. Nat Commun, 2017. 8(1): p. 126.

14. Flanagan, L.A., et al., Neurite branching on deformable substrates. Neuroreport, 2002. 13(18): p. 2411–2415.

15. Jagielska, A., et al., Mechanical environment modulates biological properties of oligodendrocyte progenitor cells. Stem Cells Dev, 2012. 21(16): p. 2905–14.

16. Moshayedi, P., et al., Mechanosensitivity of astrocytes on optimized polyacrylamide gels analyzed by quantitative morphometry. J Phys Condens Matter, 2010. 22(19): p. 194114.

17. Moshayedi, P., et al., The relationship between glial cell mechanosensitivity and foreign body reactions in the central nervous system. Biomaterials, 2014. 35(13): p. 3919–25.

18. Koser, D.E., et al., Mechanosensing is critical for axon growth in the developing brain. Nat Neurosci, 2016. 19(12): p. 1592–1598.

19. Lu, Y.B., et al., Viscoelastic properties of individual glial cells and neurons in the CNS. Proc Natl A cad Sci U S A, 2006. 103(47): p. 17759–64.

20. Koser, D.E., et al., Predicting local tissue mechanics using immunohistochemistry. bioRxiv, 2018: p. 358119.

21. Iwashita, M., et al., Systematic profiling of spatiotemporal tissue and cellular stiffness in the developing brain. Development, 2014. 141(19): p. 3793–8.

22. Sack, I., et al., Structure-sensitive elastography: on the viscoelastic powerlaw behavior of in vivo human tissue in health and disease. Soft Matter, 2013. 9(24): p. 5672–5680.

23. Millward, J.M., et al., Tissue structure and inflammatory processes shape viscoelastic properties of the mouse brain. NMR Biomed, 2015. 28(7): p. 831–9.

24. Moeendarbary, E., et al., The soft mechanical signature of glial scars in the central nervous system. Nat Commun, 2017. 8: p. 14787.

25. Eberle, D., et al., Acute but not inherited demyelination in mouse models leads to brain tissue stiffness changes. bioRxiv, 2018: p. 449603.

26. Brand, M., M. Granato, and C. Nüsslein-Volhard, Keeping and raising zebrafish. Practical approach series. Vol. 261. 2002: Oxford University Press.

27. Jung, S.H., et al., Visualization of myelination in GFP-transgenic zebrafish. Dev Dyn, 2010. 239(2): p. 592–7.

28. Wilson, J.M., R.M. Bunte, and A.J. Carty, Evaluation of Rapid Cooling and Tricaine Methanesulfonate (MS222) as Methods of Euthanasia in Zebrafish (Danio rerio). Journal of the American Association for Laboratory Animal Science, 2009. 48(6): p. 785–789.

29. Masino, M.A. and J.R. Fetcho, Fictive swimming motor patterns in wild type and mutant larval zebrafish. Journal of Neurophysiology, 2005. 93(6): p. 3177–3188.

30. Hutter, J.L. and J. Bechhoefer, Calibration of atomic-force microscope tips. Review of Scientific Instruments, 1993. 64(7): p. 1868–1873.

31. Hertz, H., Ueber die Berührung fester elastischer Körper. Journal für die reine und angewandte Mathematik, 1881. 92: p. 156–171.

32. Sneddon, I.N., The relation between load and penetration in the axisymmetric boussinesq problem for a punch of arbitrary profile. International Journal of Engineering Science, 1965. 3(1): p. 47–57.

33. Determining the elastic modulus of biological samples using the atomic force microscope. Available from: https://http://www.jpk.com/app-technotes-img/AFM/pdf/jpk-app-elastic-modulus.14-1.pdf.

34. Bolte, S. and F.P. Cordelieres, A guided tour into subcellular colocalization analysis in light microscopy. J Microsc, 2006. 224(Pt 3): p. 213–32.

35. Schindelin, J., et al., Fiji: an open-source platform for biological-image analysis. Nat Methods, 2012. 9(7): p. 676–82.

36. Scott, D.W., Multivariate Density Estimation: Theory, Practice, and Visualization. 2015: Wiley.

37. Stil, A. and P. Drapeau, Neuronal labeling patterns in the spinal cord of adult transgenic Zebrafish. Dev Neurobiol, 2016. 76(6): p. 642–60.

38. Koser, D.E., et al., CNS cell distribution and axon orientation determine local spinal cord mechanical properties. Biophys J, 2015. 108(9): p. 2137–47.

39. Christ, A.F., et al., Mechanical difference between white and gray matter in the rat cerebellum measured by scanning force microscopy. Journal of Biomechanics, 2010. 43(15): p. 2986–2992.

40. Franze, K., P.A. Janmey, and J. Guck, Mechanics in Neuronal Development and Repair. Annu Rev Biomed Eng, 2013. 15: p. 227–251.

41. Becker, T. and C.G. Becker, Regenerating descending axons preferentially reroute to the gray matter in the presence of a general macrophage/microglial reaction caudal to a spinal transection in adult zebrafish. Journal of Comparative Neurology, 2001. 433(1): p. 131–147.

42. Diekmann, H., et al., Analysis of the reticulon gene family demonstrates the absence of the neurite growth inhibitor Nogo-A in fish. Mol Biol Evol, 2005. 22(8): p. 1635–48.

43. Abdesselem, H., et al., No Nogo66- and NgR-mediated inhibition of regenerating axons in the zebrafish optic nerve. J Neurosci, 2009. 29(49): p. 15489–98.

44. Reimer, M.M., et al., Dopamine from the brain promotes spinal motor neuron generation during development and adult regeneration. Dev Cell, 2013. 25(5): p. 478–91.

45. Ballmann, C.W., et al., Stimulated Brillouin Scattering Microscopic Imaging. Sci Rep, 2015. 5: p. 18139.

46. Scarcelli, G. and S.H. Yun, In vivo Brillouin optical microscopy of the human eye. Opt Express, 2012. 20(8): p. 9197–202.

47. Schlüssler, R., et al., Mechanical Mapping of Spinal Cord Growth and Repair in Living Zebrafish Larvae by Brillouin Imaging. Biophys J, 2018. 115(5): p. 911–923.

48. Gefen, A. and S.S. Margulies, Are in vivo and in situ brain tissues mechanically similar? Journal of Biomechanics, 2004. 37(9): p. 1339–1352.

49. Weickenmeier, J., et al., Brain stiffens post mortem. Journal of the Mechanical Behavior of Biomedical Materials, 2018. 84: p. 88–98.

50. Matthews, M. and Z.M. Varga, Anesthesia and Euthanasia in Zebrafish. Ilar Journal, 2012. 53(2): p. 192–204.

51. Holtzmann, K., et al., Brain tissue stiffness is a sensitive marker for acidosis.J Neurosci Methods, 2016. 271: p. 50–4.

52. NIH. 2016 12/1/2016; Available from: https://http://www.nichd.nih.gov/health/topics/spinalinjury/conditioninfo/treatments.

