## Supplementary Material for "Zebrafish spinal cord repair is accompanied by transient tissue stiffening"

### Supplementary Methods

#### Assessment of necrosis - FDA and PI staining

Acute spinal cord sections from adult wild type fish were prepared, stored and mounted as described for indentation measurements. The tissue sections were incubated in propidium iodide (1 μg/ml in aCSF, Sigma-Aldrich, 81845) and fluorescein diacetate (5 μg/ml in aCSF, Sigma-Aldrich, F7378) for 15 min at 0.5 hours post-mortem (hpm), 2 hpm, 4 hpm, 6 hpm, 8 hpm and 72 hpm. After incubation in PI and FDA, tissue sections were washed twice in aCSF. Positive control sections were fixed with 4% PFA either immediately after vibratome sectioning or after incubation in PBS for 24 h at room temperature. Negative control sections were imaged without prior incubation in FDA and PI.

#### Assessment of apoptosis - TUNEL assay

Acute spinal cord sections from adult wild type fish were prepared and stored as described for indentation measurements. The tissue sections were immersed in 4% PFA at 0.5 hpm, 2 hpm, 4 hpm, 8 hpm and 72 hpm. To assess the amount of apoptotic cells in living tissue without the effects elicited by tissue dissection and processing, wild type zebrafish were sacrificed followed by administration of PBS and PFA through the *bulbus arteriosus* (labeled 0 hpm). After 24 hours incubation in the fixative, spinal cords were dissected, embedded, sectioned and subjected to the TUNEL assay protocol. All TUNEL assays were carried out with the *In Situ* Cell Death Detection Kit (Roche, 12156792910) according to the manufacturer's instructions. Positive control sections were digested with DNAse I (50 U/ml in 50mM Tris-HCl, pH 7.5, 1 mg/ml BSA). Negative control sections were incubated with reaction buffer without the terminal deoxynucleotidyl transferase (TdT) enzyme. In all sections, cell nuclei were additionally stained with DAPI.

**Image acquisition and processing for viability assays**

PI and FDA stained spinal cord sections were imaged using the confocal laser scanning microscope Leica TCS SP5 and a 20x/0.5 water dipping objective. Spinal cord sections subjected to the TUNEL assay were imaged by confocal fluorescence microscopy using the Zeiss LSM700 and a Plan Apochromat 10x/0.45 objective. To quantify TUNEL positive cells, the confocal stacks were subjected to the 3D cell counter plug-in implemented in Fiji. The amount of TUNEL-positive cells was normalized to the imaged volume of the confocal stack. For each time point, tissue sections were obtained from two fish and pooled for analysis. One-way analysis of variance (ANOVA) with Tukey's HSD post-hoc test using R was carried out to detect a significant influence of the *post-mortem* time interval on the number of TUNEL-positive cells. ***$p < 0.001$. $p > 0.05$ was not considered significant. Exemplary images shown were adjusted for brightness and contrast using Fiji, and assembled using CorelDraw.

**Scanning electron microscopy (SEM)**

For SEM, vibratome sections were postfixed with modified Karnovsky (2 % glutaraldehyde/2 % formaldehyde in 100 mM phosphate buffer). After several washes in PBS, the samples were postfixed in 1 % osmium tetroxide/PBS for 2 hours on ice, washed in PBS and water and dehydrated in a graded series of ethanol (30 %, 50 %, 70 %, 90 %, 96 %, 3x 100 % ethanol (pure ethanol on molecular sieve). Finally, they were critical-point dried using the Leica CPD 300 (Leica Microsystems, Vienna, Austria), mounted on 12 mm aluminium stubs and sputter-coated with gold using the Baltec SCD 050 (settings: 60 sec, 60 mA, Leica Microsystems, Vienna, Austria). Finally, the samples were analysed with a Jeol JSM 7500F cold field emission SEM (Jeol, Eching, Germany) at 8 mm working distance and 5 kV acceleration voltage using the lower secondary electron detector.

### Supplementary Information

**Acute spinal cord slices maintain viability until approximately 8 hours *post-mortem***

Despite the invasive nature of the required sample preparation, the here reported indentation measurements aim at describing intrinsic mechanical tissue properties at a near physiological state. Tissue viability and structural integrity must therefore be ensured during all sample preparation steps and measurement procedures. To assess the lifespan of acute spinal cord slices after sample preparation, and to validate the adequacy of storage and mounting conditions, a Live/Dead staining and the TUNEL method [52-55] were employed to detect necrosis and apoptosis, respectively. The Live/Dead staining used fluorescein diacetate (FDA) and propidium iodide (PI). It revealed that acute spinal cord slices displayed very strong enzymatic activity and very little, constant amount of dead cells until at least 8 hours *post-mortem* (hpm) when obtained and processed as described in this study (Suppl. Fig. 1a). Most PI-positive cells were located around the central canals, along the medial regions of the dorsal horns and at the marginal tissue regions presumably belonging to the pia mater. Fluorescein retaining, viable cells were detected in gray and white matter regions. However, white matter regions displayed very strong fluorescence intensity in comparison to gray matter. Immediate fixation in 4% paraformaldehyde (PFA) upon sectioning resulted in a large amount of dead cells and the overall loss of enzymatic activity, but no change in tissue architecture (Suppl. Fig. 1c). Incubation in PBS for 24 hours followed by PFA fixation showed that the tissue lost its structural integrity and most of its enzymatic activity indicating that the chemical composition of the immersion medium influences if the tissue architecture remains intact (Suppl. Fig. 1c). Tissue sections incubated in aCSF without FDA or PI displayed no fluorescence signal (Suppl. Fig. 1c). The TUNEL assay revealed an increase of apoptotic cells that occurred within the first 30 minutes after sacrifice, but then remained constant until at least 8 hpm before it increased further at 72 hpm (Suppl. Fig. 1b,e). The presence of apoptotic cells in immediately fixed animals (0 hpm) reflects the fact that apoptosis is a common mechanism in living, multicellular organisms to dispose of unneeded or damaged cells in a controlled manner [56]. The increase of apoptotic cells at 0.5 hpm suggests that the preparation procedure induces cell death. This could be explained by the vibratome sectioning which has been shown to produce TUNEL reactivity [57]. The sectioning induces necrosis, during which DNA fragmentation occurs, and therefore adds to the TUNEL positive population of nuclei [58-60]. Nevertheless, the constant amount of apoptotic cells until 8 hpm and the structurally intact tissue architecture as observed by microscopy indicate that the tissue remains viable and structurally preserved if kept under the aforementioned conditions.

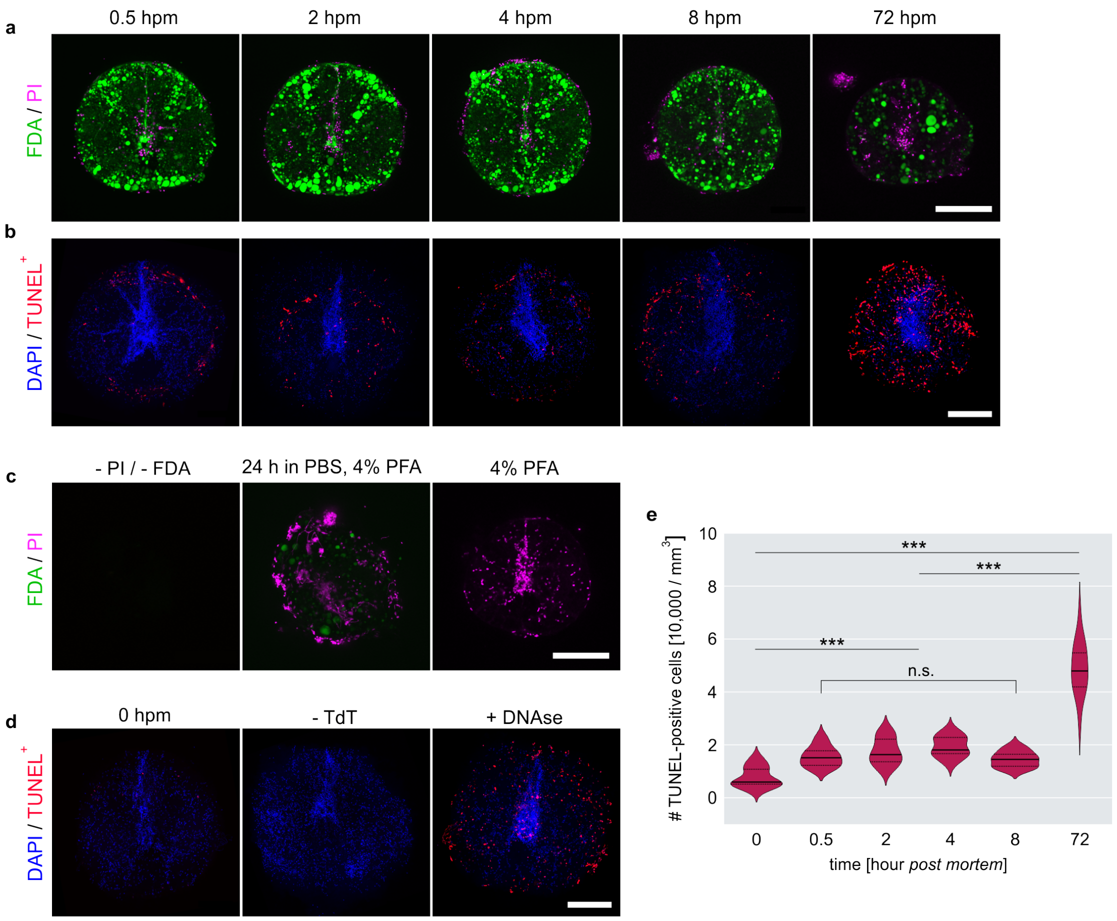


**Suppl. Fig. 1 Viability assessments of acute zebrafish spinal cord sections.** a) Assessment of necrosis using FDA (green) and PI (red) at 0.5 hours post-mortem (hpm), 2 hpm, 4 hpm, 8 hpm and 72 hpm. b) Assessment of apoptotic cells (red) using the TUNEL assay at 0.5 hpm, 2 hpm, 4 hpm, 8 hpm and 72 hpm. Nuclear counterstaining with DAPI is shown in blue. c) Control stainings for necrosis assessment without PI and FDA, after 24 h in PBS with subsequent fixation in 4% PFA and sole fixation in 4% PFA. d) Control stainings for TUNEL assay on immediately fixed cross-section (0 hpm), without TdT and after treatment with DNAse. Scale bars a) — d), 150 µm. e) Quantification of TUNEL-positive cells. ***p<0.001.

Moreover, most TUNEL-positive cells, as most PI-positive cells, were detected at the marginal regions of the tissue sections, presumably belonging to the pia mater, which could be the result of the agarose embedding. These regions were not subjected to indentation measurements as their proximity to the embedding does not allow to discern if the recorded indentation response originates from the spinal cord’s material properties, that of the agarose or a combination of both. The results obtained from both necrosis and apoptosis assessments show that the sample preparation steps as well as the storage and mounting conditions do not impair the overall tissue viability or tissue architecture significantly up until 8 hpm. However, to account for seasonal changes of temperature and intraspecies variations, all experiments using acute zebrafish spinal cord sections were restricted to a 5 hpm time interval. The results obtained from the viability assays indicate that indentation measurements carried out with acute zebrafish spinal cord sections *ex vivo* appear to be suitable to characterize mechanical properties of the spinal cord tissue close to *in vivo* conditions.

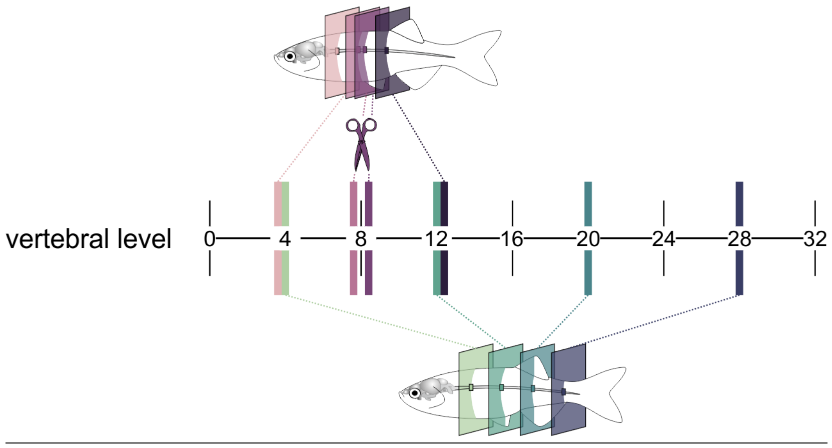


**Suppl. Fig. 2 Overview of vertebral levels.** Spinal cord sections from uninjured animals were obtained from the levels of the 4th, the 8th, the 20th and the 28th vertebra. Spinal cord sections from spinal cord transected and sham-operated animals were obtained from vertebral levels that correspond to the 4th, the 7-8th (V8-) , 8-9th (V8+) and 12th vertebra. Sham-operated animals additionally provided a tissue section at the level of the 8th vertebra, where the injury site and glial bridge in spinal cord transected animals is located.

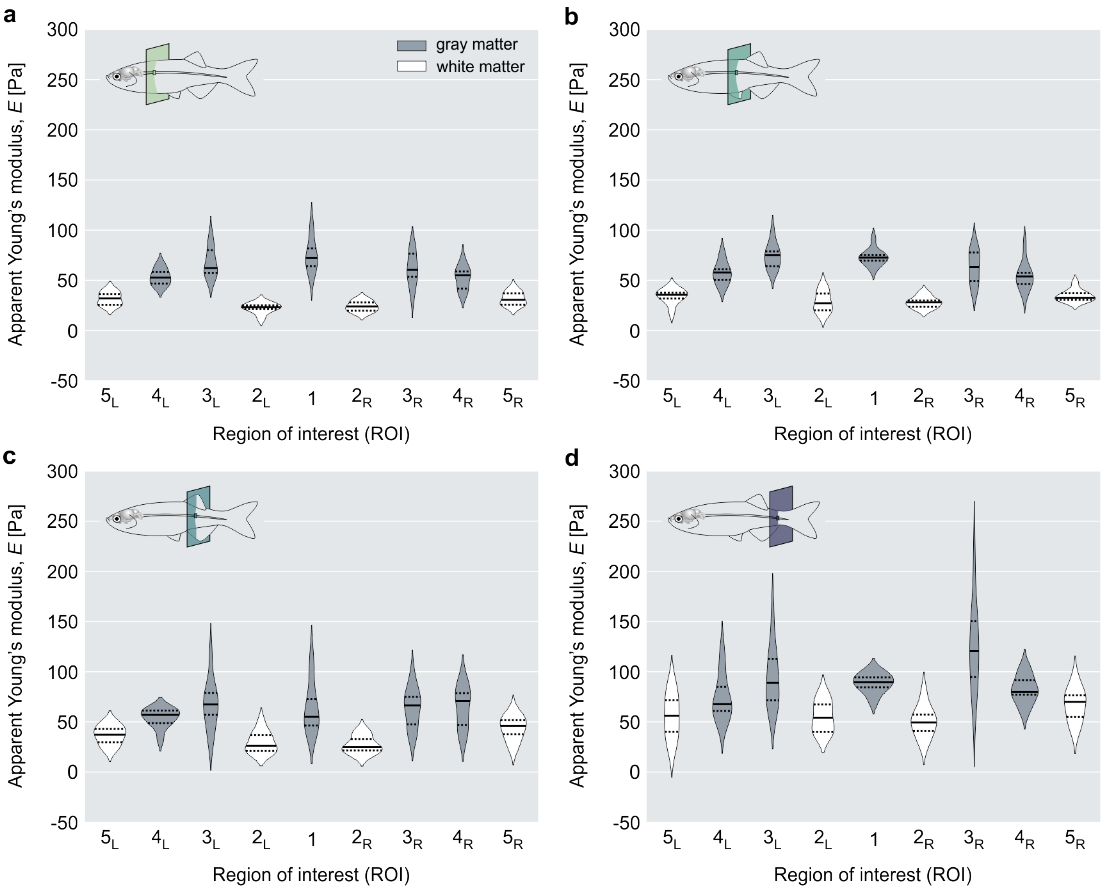


**Suppl. Fig. 3 Distribution of apparent Young's moduli of individual ROIs along the anterior-posterior axis in transgenic zebrafish at 6 mpf.** Spinal cord tissue sections were obtained from uninjured fish ($n$ = 10) from a) vertebral level V4, b) vertebral level V12, c) vertebral level V20 and d) vertebral level V28.

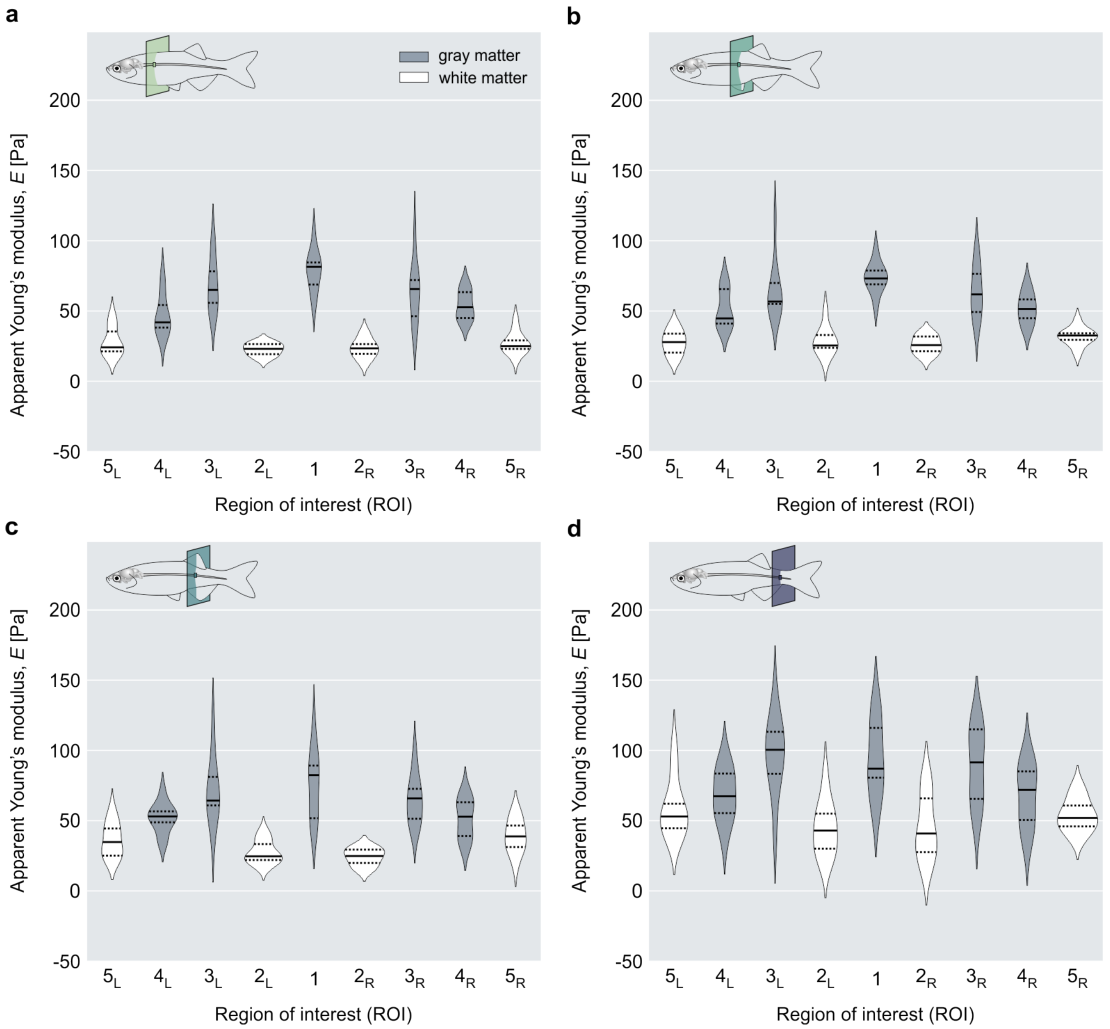


**Suppl. Fig. 4 Distribution of apparent Young's moduli of individual ROIs along the anterior-posterior axis in transgenic zebrafish at 9 mpf.** Spinal cord tissue sections were obtained from uninjured fish ($n$ = 10) from a) vertebral level V4, b) vertebral level V12, c) vertebral level V20 and d) vertebral level V28.

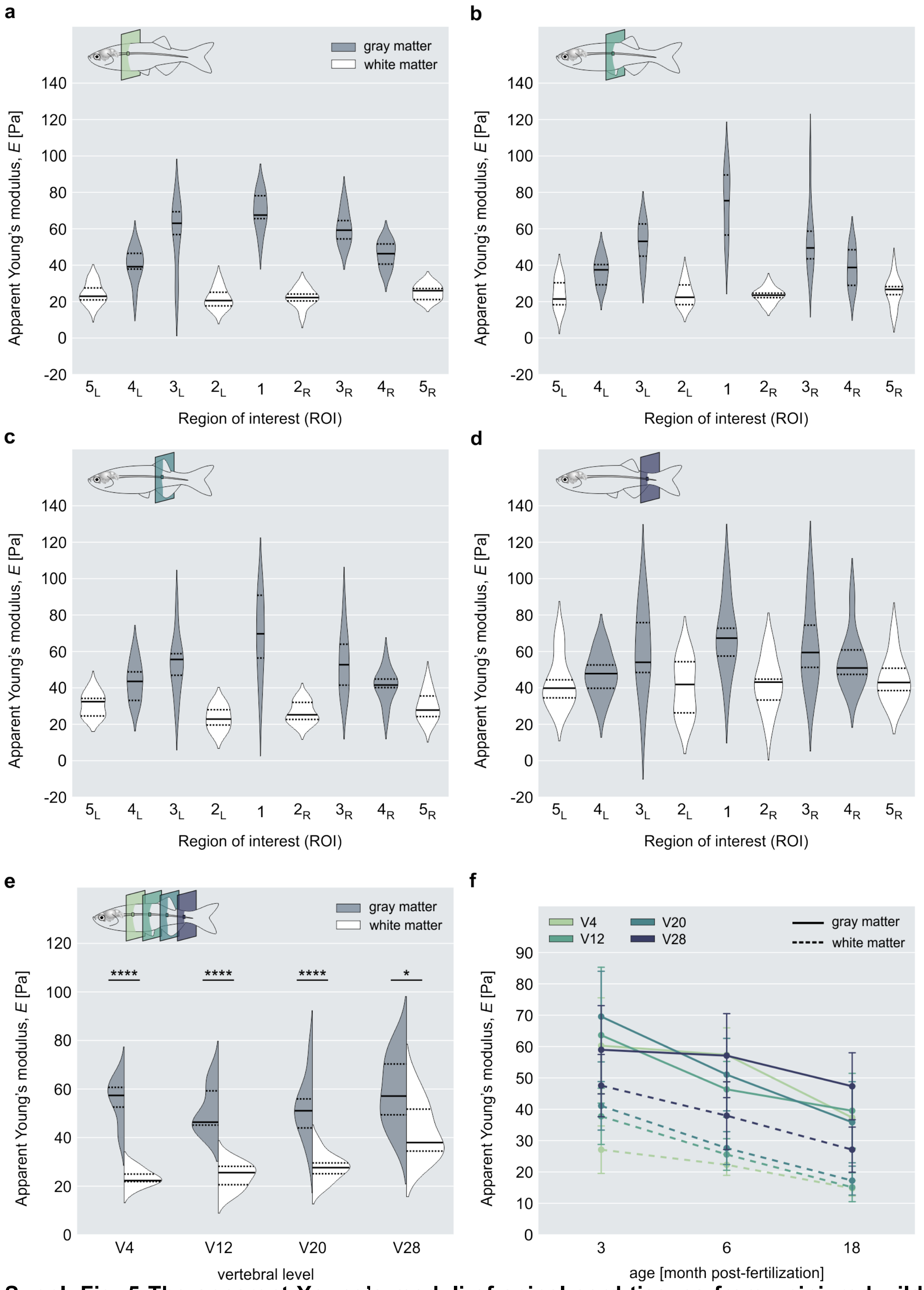


**Suppl. Fig. 5 The apparent Young's moduli of spinal cord tissues from uninjured wild type zebrafish.** a) – d) Distribution of apparent Young's moduli of individual ROIs along the anterior-posterior axis in wild type zebrafish at 6 mpf . Spinal cord tissue sections were obtained from uninjured fish (*n* = 10) from a) vertebral level V4, b) vertebral level V12, c) vertebral level V20 and d) vertebral level V28. e) The distribution of combined gray and white matter at the indicated vertebral levels at 6 mpf. Solid lines and dashed lines represent medians and interquartile ranges, respectively. $*p<0.05$, $****p < 0.0001$. f) Comparison of the apparent Young's moduli of spinal gray and white matter at different vertebral levels (V4-V28) at 3 months post-fertilization (mpf) (*n* = 7), 6 mpf (*n* = 9) and 18 mpf (*n* = 6). Solid lines and dashed lines represent gray and white matter, respectively.

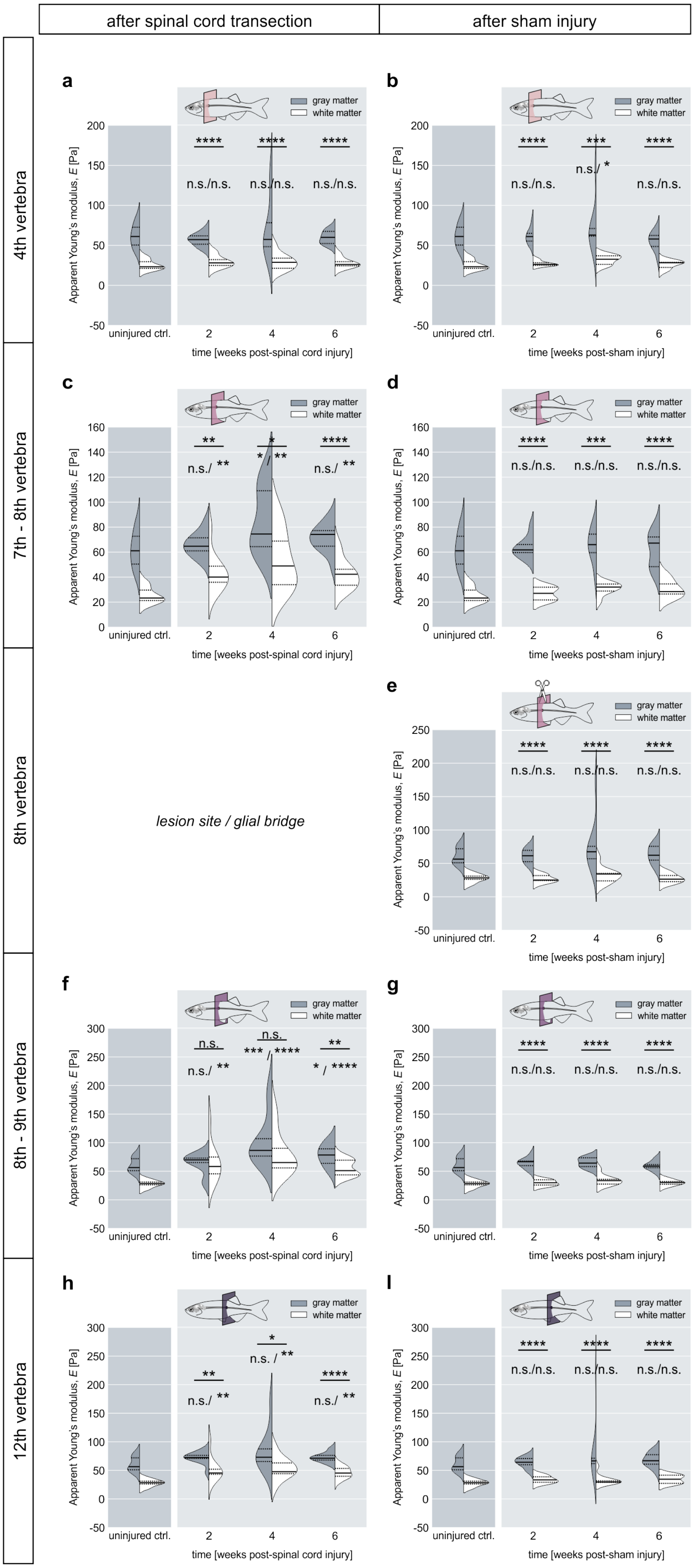

after spinal cord transection
after sham injury
4th vertebra
7th - 8th vertebra
8th vertebra
8th - 9th vertebra
12th vertebra
a
b
c
d
e
f
g
h
I
gray matter
white matter
Apparent Young's modulus, E [Pa]
uninjured ctrl.
2
4
6
time [weeks post-spinal cord injury]
time [weeks post-sham injury]
****
***
**
*
n.s./n.s.
n.s./ *
n.s./ **
* / **
n.s.
*** / ****
* / ****
n.s. / **
lesion site / glial bridge

**Suppl. Fig. 6 Overview of the spatio-temporal evolution of apparent Young's moduli of transgenic zebrafish after spinal cord injury and after sham injury.** Regenerating spinal cords were subjected to mechanical testing at 2 wpi (*n* = 10), 4 wpi (*n* = 11) and 6 wpi (*n* = 13). Spinal cords from sham-operated fish were probed at the same time points 2 wpi (*n* = 10), 4 wpi (*n* = 10) and 6 wpi (*n* = 10). Significance levels correspond to pairwise comparisons between combined gray and white matter and to pairwise comparisons with gray and white matter from uninjured control animals (*n* = 10), respectively. Tissue sections from uninjured control animals were located at the 4$^{th}$ vertebral level and compared to tissue sections located rostrally to the lesion site. Tissue sections caudally located to the lesion site were compared to uninjured control sections from the 12$^{th}$ vertebral level (see Suppl.Fig.2). *p<0.05, **p<0.01, ***p<0.001, ****p < 0.0001.

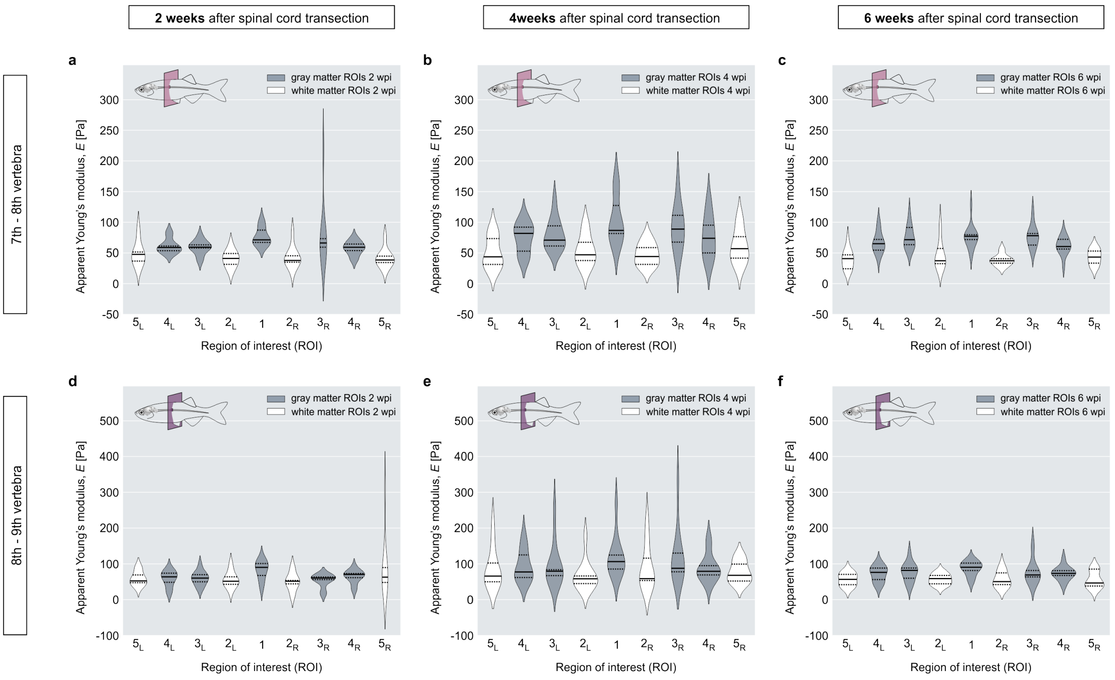


**Suppl. Fig. 7 The apparent Young's moduli of individual ROIs after spinal cord injury in transgenic zebrafish.** Tissue sections were located proximal to the lesion site and probed at 2 wpi (*n* = 10), 4 wpi (*n* = 11) and 6 wpi (*n* = 13).

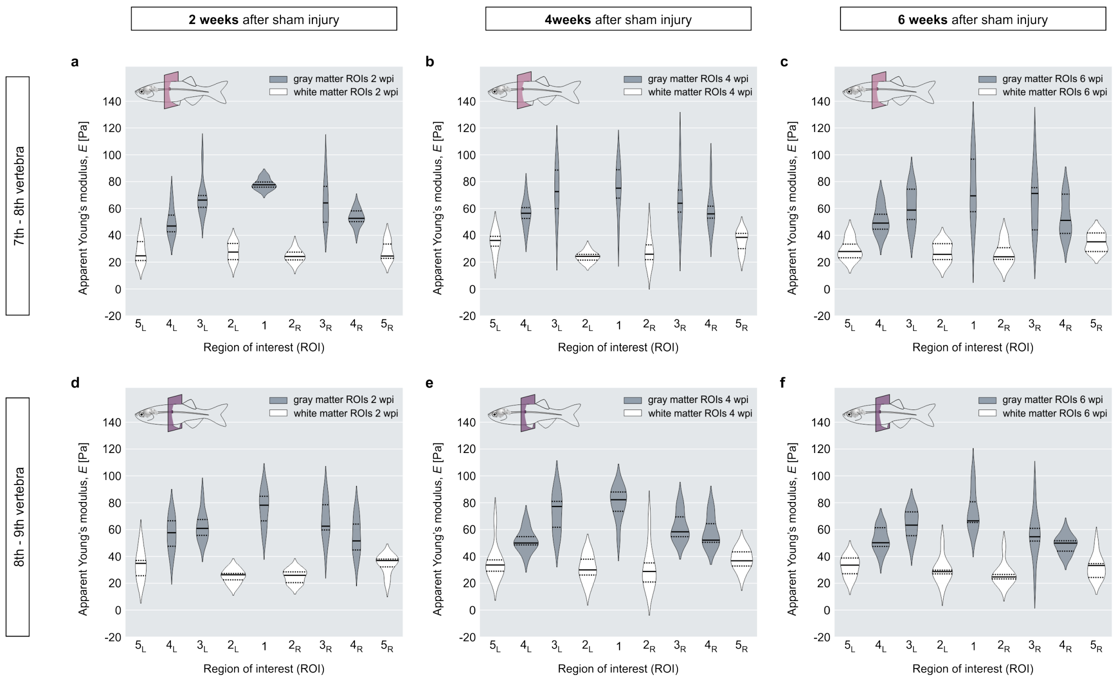


**Suppl. Fig. 8 The apparent Young's moduli of individual ROIs after sham injury in transgenic zebrafish.** Tissue sections were located proximal to the lesion site and probed at 2 wpi (*n* = 10), 4 wpi (*n* = 10) and 6 wpi (*n* = 10).

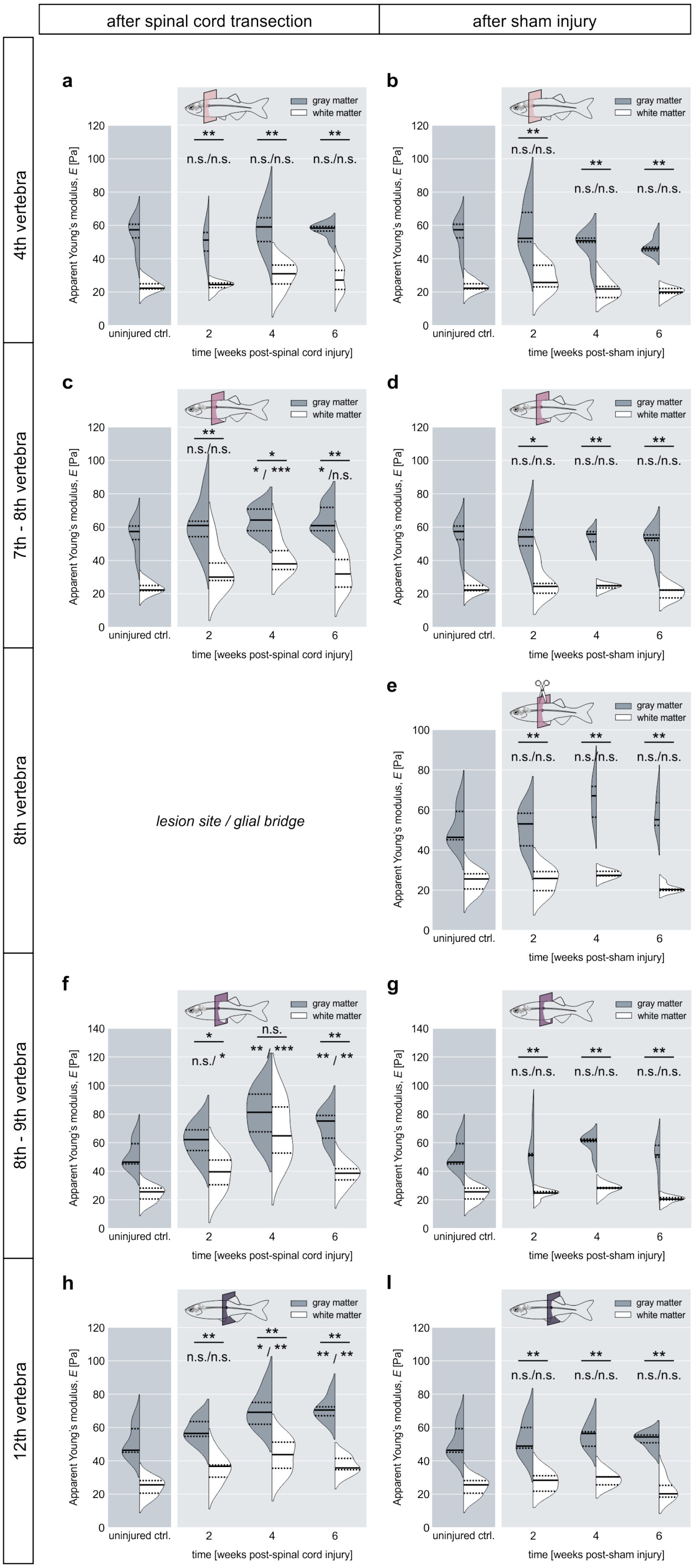

after spinal cord transection
after sham injury
4th vertebra
7th - 8th vertebra
8th vertebra
8th - 9th vertebra
12th vertebra
a
b
c
d
e
f
g
h
l
gray matter
white matter
Apparent Young's modulus, E [Pa]
uninjured ctrl.
time [weeks post-spinal cord injury]
time [weeks post-sham injury]
lesion site / glial bridge
n.s./n.s.
* / ***
* /n.s.
n.s./ *
** / ***
** / **
* / **

**Suppl. Fig. 9 Overview of spatio-temporal evolution of apparent Young's moduli of wild type zebrafish after spinal cord injury and sham injury.** Regenerating spinal cords were subjected to mechanical testing at 2 wpi ($n$ = 6), 4 wpi ($n$ = 6) and 6 wpi ($n$ = 6). Spinal cords from sham-operated fish were probed at the same time points 2 wpi ($n$ = 5), 4 wpi ($n$ = 5) and 6 wpi ($n$ = 5). Significance levels correspond to pairwise comparisons between combined gray and white matter and to pairwise comparisons with gray and white matter from uninjured control animals ($n$ = 10), respectively. Tissue sections from uninjured control animals were located at the 4$^{th}$ vertebral level and compared to tissue sections located rostrally to the lesion site. Tissue sections caudally located to the lesion site were compared to uninjured control sections from the 12$^{th}$ vertebral level (see Suppl.Fig. 2). *$p<0.05$, **$p<0.01$, ***$p<0.001$, ****$p < 0.0001$.

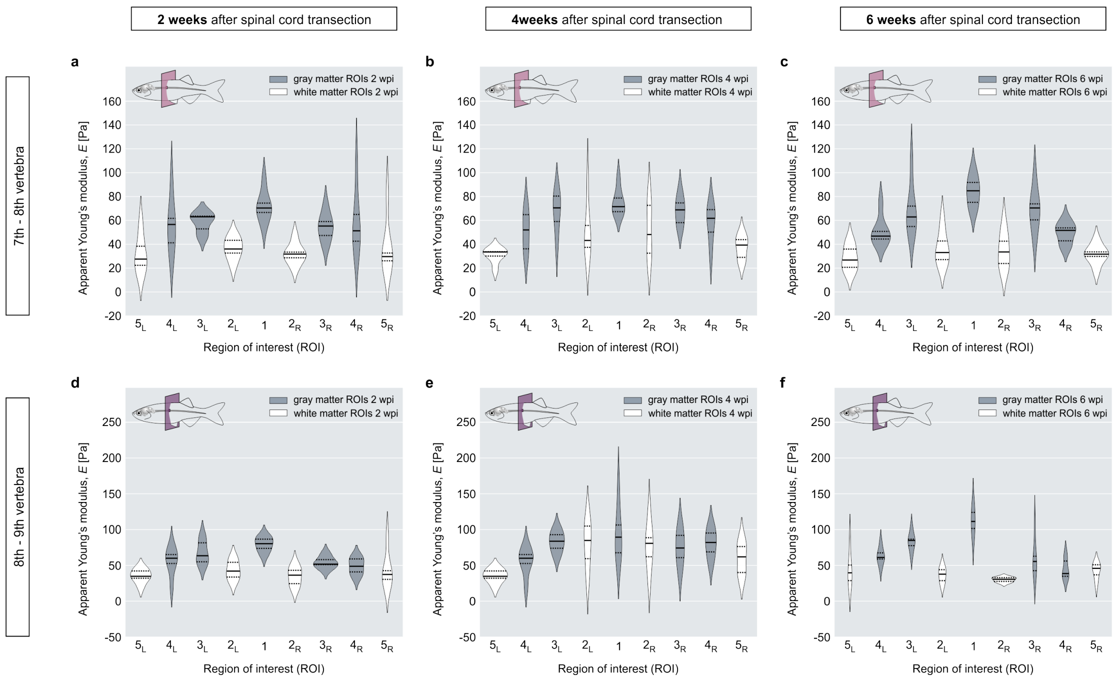


**Suppl. Fig. 10 The apparent Young's moduli of individual ROIs after spinal cord injury in wild type zebrafish.** Tissue sections were located proximal to the lesion site and probed at 2 wpi (*n* = 6), 4 wpi (*n* = 6) and 6 wpi (*n* = 6).

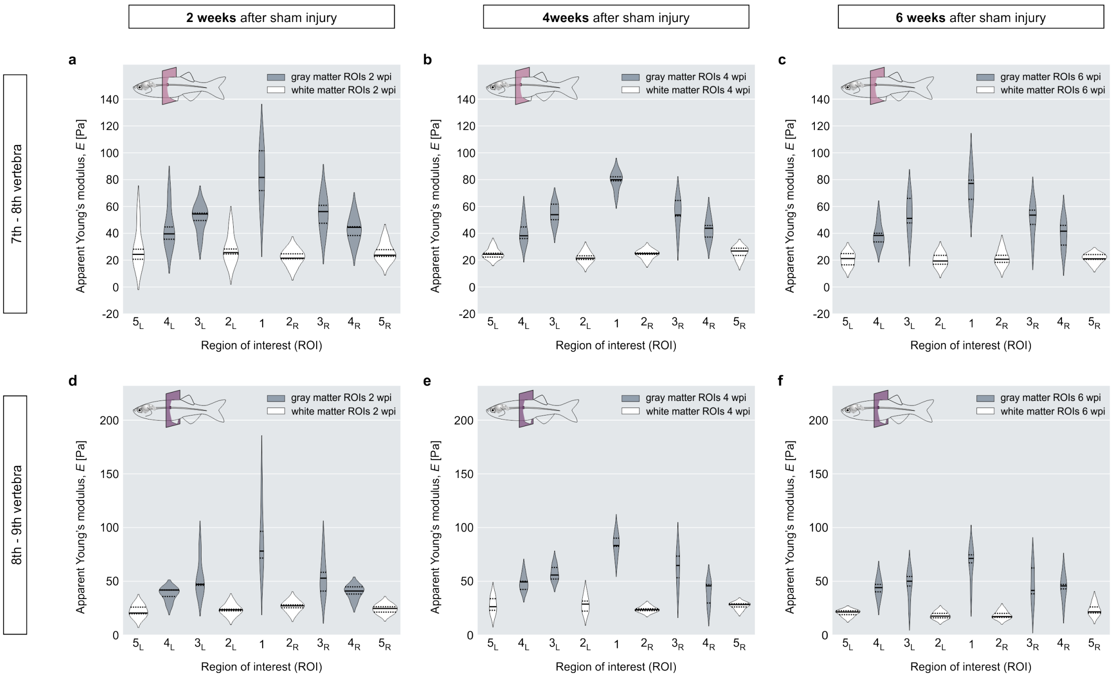


**Suppl. Fig. 11 The apparent Young's moduli of individual ROIs after sham injury in wild type zebrafish.** Tissue sections were located proximal to the lesion site and probed at 2 wpi (*n* = 5), 4 wpi (*n* = 5) and 6 wpi (*n* = 5).

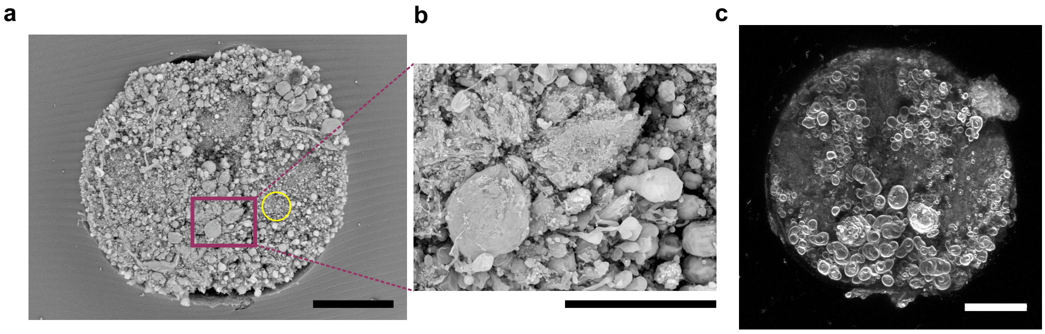


**Suppl. Fig. 12 Surface characterization of zebrafish spinal cord sections.** a) Scanning electron microscopy (SEM) image of a spinal cord section that was subjected to the preparation procedures used for indentation measurements prior to SEM preparations. The yellow circle represents the diameter of the indenter. Scale bar, 100 µm. b) High-resolution scan of indicated area in a). Scale bar, 30 µm. c) An acutely prepared spinal cord section was stained with FM-143 which provides a non-specific staining of cell-surface membranes. The tissue section was imaged as described for tissue sections subjected to viability assays using FDA and PI. Scale bar, 100 µm.
